# Cooperativity and induced oligomerisation control the interaction of SARS-CoV-2 with its cellular receptor and patient-derived antibodies

**DOI:** 10.1101/2023.09.14.557399

**Authors:** Roi Asor, Anna Olerinyova, Sean A. Burnap, Manish S. Kushwah, Fabian Soltermann, Lucas Powell Rudden, Mario Hensen, Snežana Vasiljevic, Juliane Brun, Michelle Hill, Liu Chang, Wanwisa Dejnirattisa, Piyada Supasa, Juthatip Mogkolsapaya, Daming Zhou, David I. Stuart, Gavin R. Screaton, Matteo Degiacomi, Nicole Zitzmann, Justin L. P. Benesch, Weston B. Struwe, Philipp Kukura

## Abstract

Viral entry is mediated by oligomeric proteins on the virus and cell surfaces. The association is therefore open to multivalent interactions between these proteins, yet such recognition is typically rationalised as affinity between monomeric equivalents. As a result, assessment of the thermodynamic mechanisms that control viral entry has been limited. Here, we use mass photometry to overcome the analytical challenges consequent to multivalency. Examining the interaction between the spike protein of SARS-CoV**-**2 and the ACE2 receptor, we find that ACE2 induces oligomerisation of spike in a variant-dependent fashion. We also demonstrate that patient-derived antibodies use induced-oligomerisation as a primary inhibition mechanism or to enhance the effects of receptor-site blocking. Our results reveal that naive affinity measurements are poor predictors of potency, and introduce a novel antibody-based inhibition mechanism for oligomeric targets.

**One-Sentence Summary:** Multivalent interactions between viral proteins, cell-surface receptors, and anti-viral antibodies regulate infection and inhibition.

## Introduction

Successful viral entry requires efficient engagement of receptors on the host cell by proteins on the virus surface (*1*). Such interactions are targeted both by some aspects of the host immune system and various anti-viral therapeutics, in an effort to out-compete the native association. Most proteins on the virus and cell surfaces are oligomeric, and therefore the underlying molecular interactions between them offer great scope for both the virus and immune system to leverage the thermodynamic benefits of multivalent binding (*2-6*). While there is a growing appreciation of the importance of multivalency, over and above the strength of the constituent interfaces, its visualisation and quantification are challenging due to the inevitable heterogeneity of assembly states it gives rise to.

SARS-CoV-2 infection and inhibition represent an archetypal system for studying multivalency: it involves a trimeric viral envelope fusion spike protein that attaches to a dimeric angiotensin-converting enzyme 2 (ACE2) on host cells (**Fig. 1a**) (*7*). It is intuitive to assume, and frequently stated, that infectivity of SARS-CoV-2 variants correlates with the affinity of their form of spike to ACE2 (*8-11*). This correlation, however, is not universal and numerous reports have reported weaker affinities for more infectious variants, instead invoking improved antibody evasion to explain increased infectivity (*12-14*).

**Figure 1.**
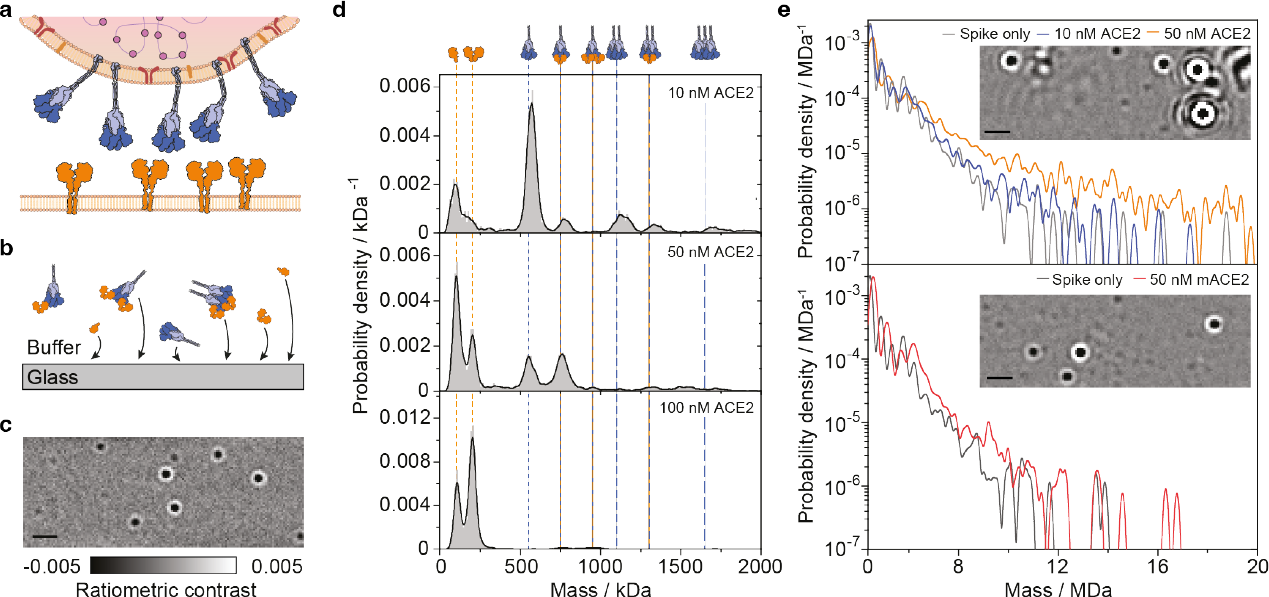
ACE2 induces oligomerisation of spike trimers in solution. (**a**) Schematic of the multivalent interaction partners at the SARS-CoV2 virus-cell interface, containing viral trimeric spike glycoproteins and dimeric ACE2 on the surface of the host cell. (**b**) The detection principle of solution-based mass photometry, relying on non-specific binding of soluble proteins to a glass surface. (**c**) Resulting MP images of individual complexes from a spike-ACE2 mixture. Scale bar: 1μm. (**d**) Mass histograms of spike-ACE2 mixtures. Spike trimers at 0.55 μM were incubated in the presence of 0.33 to 3.3 μM ACE2 for 10 min at ambient room temperature, and then rapidly diluted to the working concentration of MP just before data acquisition. The final concentration of spike trimer was 16.7 nM. Vertical lines indicate the masses of the expected molecular complexes. Histograms were generated by combining 3 -6 technical replicates. The total number of particles were 12430, 28246 and 86390 for samples containing 10, 50, and 100 nM ACE2, respectively. (**e**) Mass histograms, for mixing spike trimers with ACE2 (top) and monomeric ACE2 (mACE2, bottom), presented on a semi logarithmic scale, showing the increase in the solution concentration of large spike-ACE2 complexes. The probability density was calculated using kernel width of 100 kDa and included all particles larger than 450 kDa to exclude the varying contribution of free ACE2. Insets show representative frames of the recorded MP video of spike with 50 nM ACE2 and 50 nM mACE. Scale bar: 1 μM.

In terms of inhibition, ACE2 has emerged as an early potential therapeutic (*15*), that exhibits strongly enhanced binding to spike and a much lower IC50 compared to a form of ACE2 that cannot dimerise (*11, 16, 17*). These results point towards avidity effects: the dimerization of ACE2 results in tighter binding than can be explained by simple combination of two monomers. Essentially equivalent observations have been made in the context of antibody efficacy, where stark differences have been observed between intact IgG antibodies and single Fabs (*5*). As for ACE2, multivalency is an important consideration for antibodies, given that they are composed of an Fc region fused to two identical Fabs effectively acting as covalent dimers.

Despite this strong evidence for the involvement and significant influence of multivalency on both SARS-CoV-2 host-cell binding (*6*) and inhibition of viral infection by antibodies more broadly (*3*), insight into its strength, and how it depends on different virus variants and antibody identity, is absent. The reason for this is that structural and biophysical characterization often relies on simplifying the system by reduction to monomeric interactions. For instance, structures relevant to the spike-ACE2 interaction have been solved largely for scenarios where one of the binding partners is engineered to be monomeric, while quantification of affinities is interpreted within a simple 1:1 binding model, such that any avidity effects are hidden within an apparent enhancement of the observed binding affinity. As a result, despite a tremendous number of studies emerging over the past few years aimed at understanding the connection between affinity, infectivity and inhibition for SARS-CoV2, no clear picture has emerged.

To address these shortcomings, we designed an approach based on mass photometry (MP) (*18*) that enables us to observe and quantify receptor-ligand interactions at the molecular level, both free in solution and confined to lipid bilayers (*19, 20*) that mimic the surface membrane of the virus and host cells. The mass resolution and single-molecule sensitivity of MP enables digital counting of individual proteins and their complexes, which can then be separated into affinities for each elementary step within the coupled equilibria that comprise the heterogeneous system (*21*), rather than aggregating them into a 1:1 binding model. We find that induced-oligomerisation and cooperativity can be directly observed and quantified from the resulting mass distributions for spike and ACE2 or antibodies. This allows us to pinpoint the precise molecular steps that are leveraged by the virus and immune system for efficient association with the cell, or its inhibition, revealing a central role of induced oligomerisation for both cellular binding and antibody-based inhibition.

## Results

We begin by characterising the interactions that occur in solution, for which we use a standard MP assay, where protein complexes binding non-specifically from solution to a glass surface are mass-measured from their images (**Fig. 1b,c**). Quantifying the contrast of each binding event together with a mass calibration results in mass distributions representative of protein complexes present in solution. For our Wuhan-Hu-1 spike construct (wtSpike), we find a major species near the expected 550 kDa trimer mass, as well as additional peaks corresponding to higher oligomers of the trimer (**SI Fig. 1a**). Performing such experiments for mixtures of ACE2 and RBD yields the full interaction landscape, with a binding affinity of 28 ± 9 nM between a single RBD and individual ACE2 binding site (**SI Fig. 2**), in excellent agreement with previous results (*7, 8, 11, 12, 22-24*). These results demonstrate the validity of our method to identify different oligomeric species, their binding to ligands and thus to quantify affinities with stoichiometric resolution.

**Figure 2.**
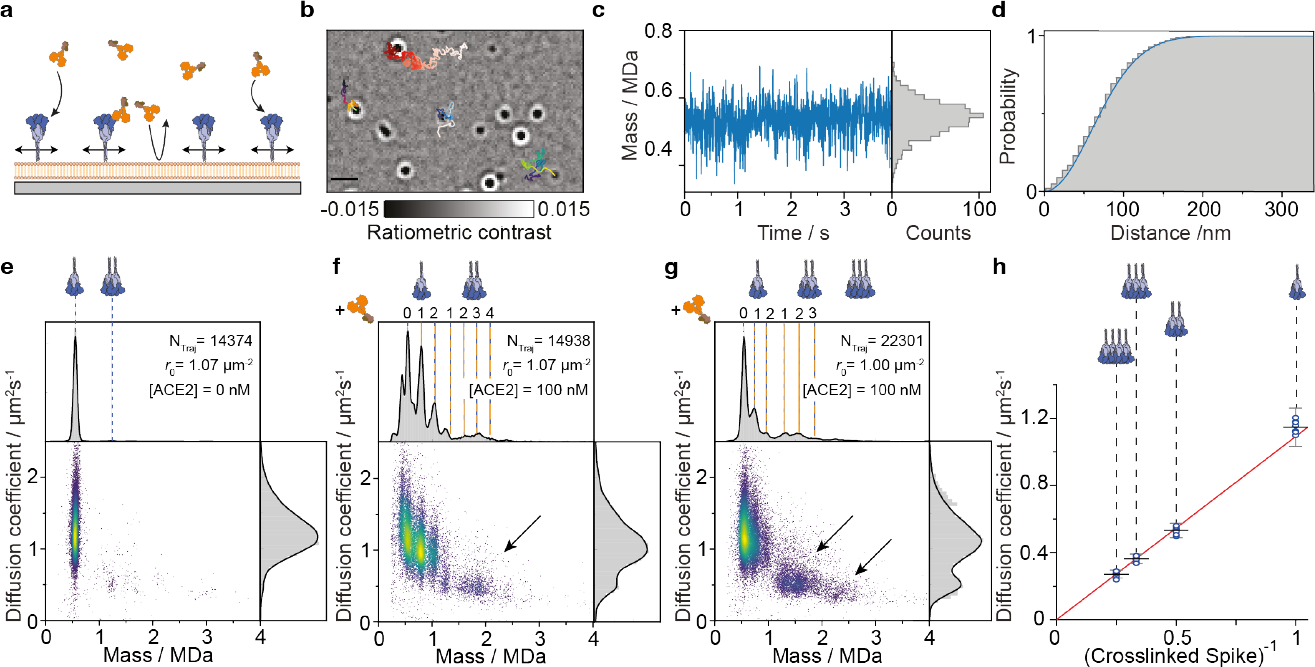
Single particle tracking and mass measurement reveal ACE2-induced oligomerisation of spike trimers on supported lipid bilayers. (**a**) Schematic of the assay with bilayer-tethered spike trimers and ACE2 binding from solution. The supported lipid bilayers also serve as a passivation layer keeping the concentration of ACE2 constant in solution. (**b**) Representative frame from a median ratiometric video, including trajectories for a few particles. The colour gradient corresponds to trajectory propagation time.(**c**) Representative recorded single-particle mass trace, and the corresponding mass histogram of an individual spike trimer measured at 270 Hz (grey). The black curve corresponds to the intrinsic measurement noise distribution in mass units. The distribution was shifted to be centered at the average particle mass. (**d**) Cumulative probability distribution of the distance travelled by a single spike trimer during a single frame within its measured trajectory. The corresponding time interval for particle displacement was 3.7 ms. The blue curve corresponds to the best fitted model used to extract the diffusion coefficient (see **SI**). (**e-g**) Two dimensional plots of the measured diffusion coefficient *vs*. average mass of individual trajectories (scatter points) for (**e**) wtSpike following equilibration for a few hours on the supported lipid bilayer, (**f**) wtSpike and (**g**) omSpike, both following equilibration with 100 nM ACE2. The expected masses for different stoichiometries are indicated by vertical lines. The number of trajectories recorded for each condition (*N*_traj_) and detected initial surface density of spike (*ρ*_0_) are stataed in each panel. Arrows indicate the trajectories that correspond to complexes of crosslinked Spike trimers. (**h**) Measured diffusion coefficient *vs*. the inverse of Spike cluster size for tethered wtSpike. Circles indicate the diffusion coefficient estimate based on a two-dimensional Gaussian mixture model from eight independent experiments (**SI Fig. 10**), black bars and error bars correspond to the averages and their standard deviations. Error bars for individual circles are not shown for clarity. The red line corresponds to a linear fit with the intercept set to the origin.

**Figure 3.**
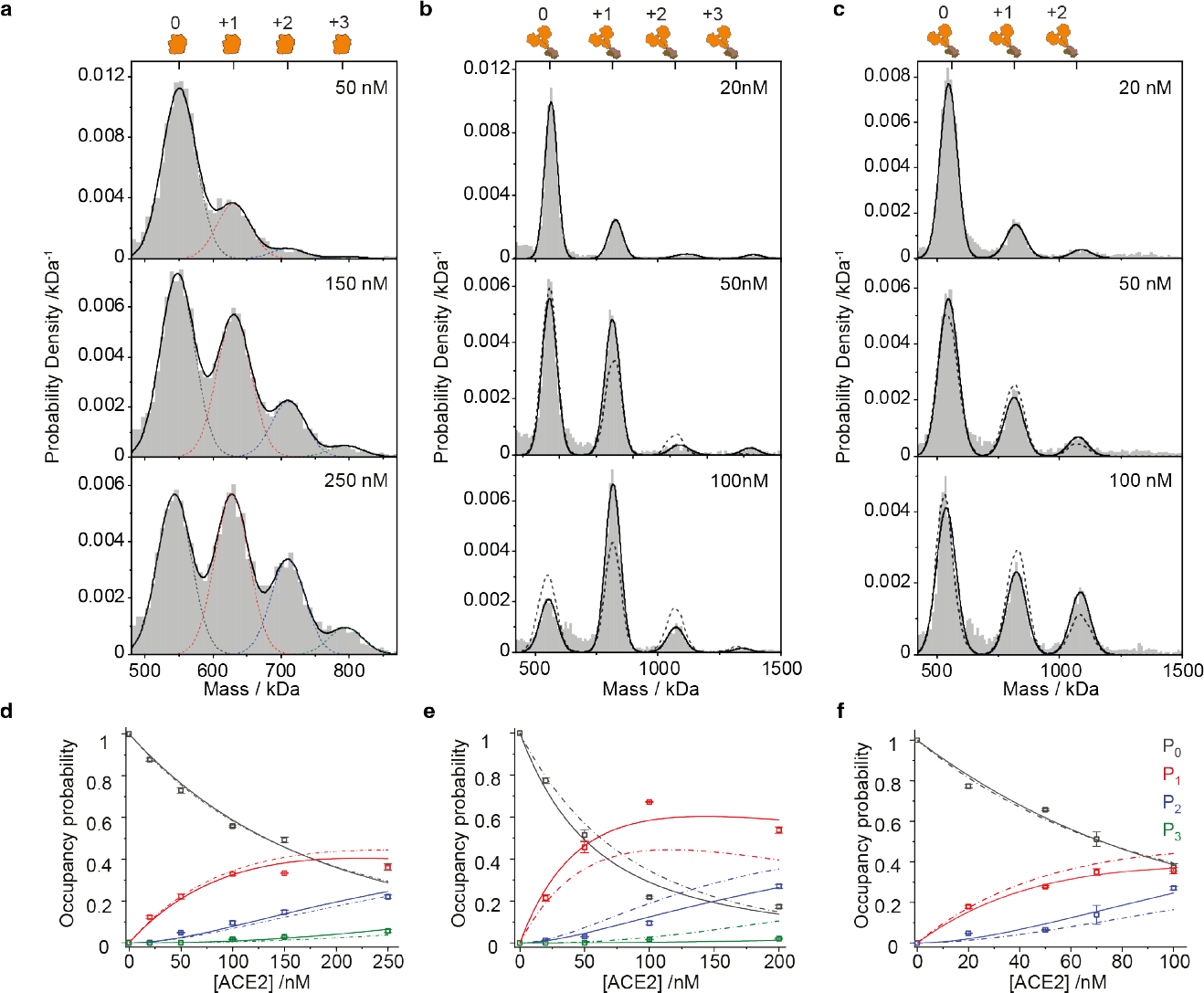
Thermodynamic analysis of the Spike-ACE2 interaction in solution reveals variant specific cooperativity. (**a-c**) Normalised mass distributions for mixtures of 25 nM wtSpike with increasing concentrations of (**a**) mACE2 and (**b**) ACE2, and 25 nM omSpike with increasing concentrations of ACE2 (c). Histograms represent cumulative counts of 3-5 technical repeats per mixture. The black curve corresponds to the modelled distribution based on a fitted sum of Gaussian functions (black broken lines). Gaussian functions were constrained to the expected masses of the individual complexes and to their expected experimental mass standard deviation. The expected positions of the different spike:ACE2 stoichiometries are indicated at the top of each panel. (**d-f**) The resolved occupancy probabilities (scatter points) based on the fitted Gaussian functions for spike with 0 (free spike), 1, 2 and 3 bound (**d**) mACE2,(**e**) ACE2 to wtSPike and (**f**) ACE2 to omSpike as a function of ACE2 concentration. Individual scatter symbols and error bars correspond to the average values of 3-5 technical replicates and their standard deviations, respectively. Solid lines correspond to the expected occupancies based on the globally best fitted thermodynamic model. The model takes into account a fundamental standard free energy change for interaction between individual RBD site and ACE2 monomer, Δ*G*°, the degeneracy of the multivalent subunits, and an effective free energy term to account for cooperativity between RBDs on the same Spike trimer, δΔ*G*° (**see Section 2 in SI**). Broken lines represent the best fitted model, assuming no cooperativity. For omSpike (**f**) we assumed a maximum coverage of 2 ACE2 given no evidence for three bound ACE2 on a single spike trimer. Dashed lines in panels **b** and **c** (50 and 100 nM ACE2) show the expected shape of the mass distribution in the absence of cooperativity (δΔ*G*° = 0).

### ACE2 induces oligomerization of wtSpike in solution

Equipped with these capabilities, we turned to quantify the interaction between the oligomeric binding partners, namely the solubilised versions of wtSpike and ACE2. Previous studies have reported extremely tight binding affinities on the order of few nM or less (*11, 13, 25, 26*). Measurements of 50 nM wtSpike at 1:1.67 and 3:1 ACE2 to wtSpike trimer ratios after incubation at μM concentration revealed binding of 1 and 2 ACE2 to the wtSpike trimer, and oligomers thereof (**Fig. 1d** and **SI Fig. 1**). Interestingly, under both conditions, we found clear signatures of free ACE2 and spike trimer, pointing towards a much weaker binding affinity (>40 nM) compared to those previously reported. The changeover from predominantly monomeric to dimeric ACE2 in solution over the explored concentration range is expected based on the our measured *K*_D_ (12 ± 2 nM, **SI Fig. 2 and SI Table 1**).

At a 6:1 ACE2:wtSpike trimer ratio, we observed a dramatic decrease in the number of detected spike-containing species in the <2 MDa mass range, indicating almost complete loss of wtSpike from solution. Inspection of the resulting MP images revealed signatures exhibiting very large optical contrast that must stem from more massive particles than those generated by wtSpike trimers alone (SI Movie 1, **Fig. 1e**, inset). These are very low in abundance, so the single-particle sensitivity of MP is highly advantageous because it provides in principle unlimited dynamic range for detection and quantification, including of large complexes. Indeed, when plotting the mass distributions from these experiments on a logarithmic abundance scale, we find a long tail of large oligomers with masses >4 MDa that increases with increasing ACE2 to wtSpike ratio persisting all the way to 20 MDa (**Fig. 1e** and **SI Fig. 1b**,**c and 3**).

To evaluate if the occurrence of these large objects arises from induced oligomerisaton of wtSpike by ACE2, we repeated these experiments under similar conditions with mACE2. The resulting MP images now resembled those for wtSpike-only (**Fig. 1e**, inset), without any significant increase in the abundance of species of mass >4 MDa. Taken together, these results suggest that ACE2 induces oligomerisation of wtSpike in solution. They may also help explain the lack of structural studies using spike trimers and ACE2, because this combination leads to a heterogeneous mixture that is not suitable to single particle averaging required for cryo-electron microscopy. In addition, induced oligomerization explains the discrepancy between our observation of a relatively weak ACE2-wtSpike interaction (>40 nM) and previous reports (<3 nM) obtained with traditional assays. This is likely due to the fact that such surface-based assays use a dense layer of one of the ligands, which is intrinsically subject to avidity effects when interacting with oligomeric ligands from solution. Our results call for a re-evaluation of both the interaction strength between spike and ACE2, as well as more generally questioning the suitability of standard surface-based assays when using oligomeric interaction partners, importantly including full-length antibodies.

### Induced oligomerisation of spike by ACE2 proceeds on bilayer membranes

These experiments demonstrate that ACE2 induces oligomerisation of spike in solution, but cannot reveal the associated energetics, nor does it resemble the membrane-associated nature of the interacting species. To address this, we turned to an alternative approach to MP, based on tracking individual proteins diffusing on supported lipid bilayers (*19, 20*). We attach individual spike trimers to lipids in the bilayer membrane using a 6xHis tag-NTA(Ni) linkage (**Fig. 2a**), and then follow their motion and mass as a function of time (**Fig. 2b**, SI Video 2). In this way, we can monitor individual spike trimers for extended periods of time, measuring their mass continuously on a frame-by-frame basis (**Fig. 2c**). The observed mass fluctuations are a consequence of the high frame rate required to minimise motional blurring (270 Hz) as seen by the overlay of the measurement noise distribution (**Fig. 2c**), rather than actual mass fluctuations of the complexes (**SI Fig. 4**).

**Figure 4.**
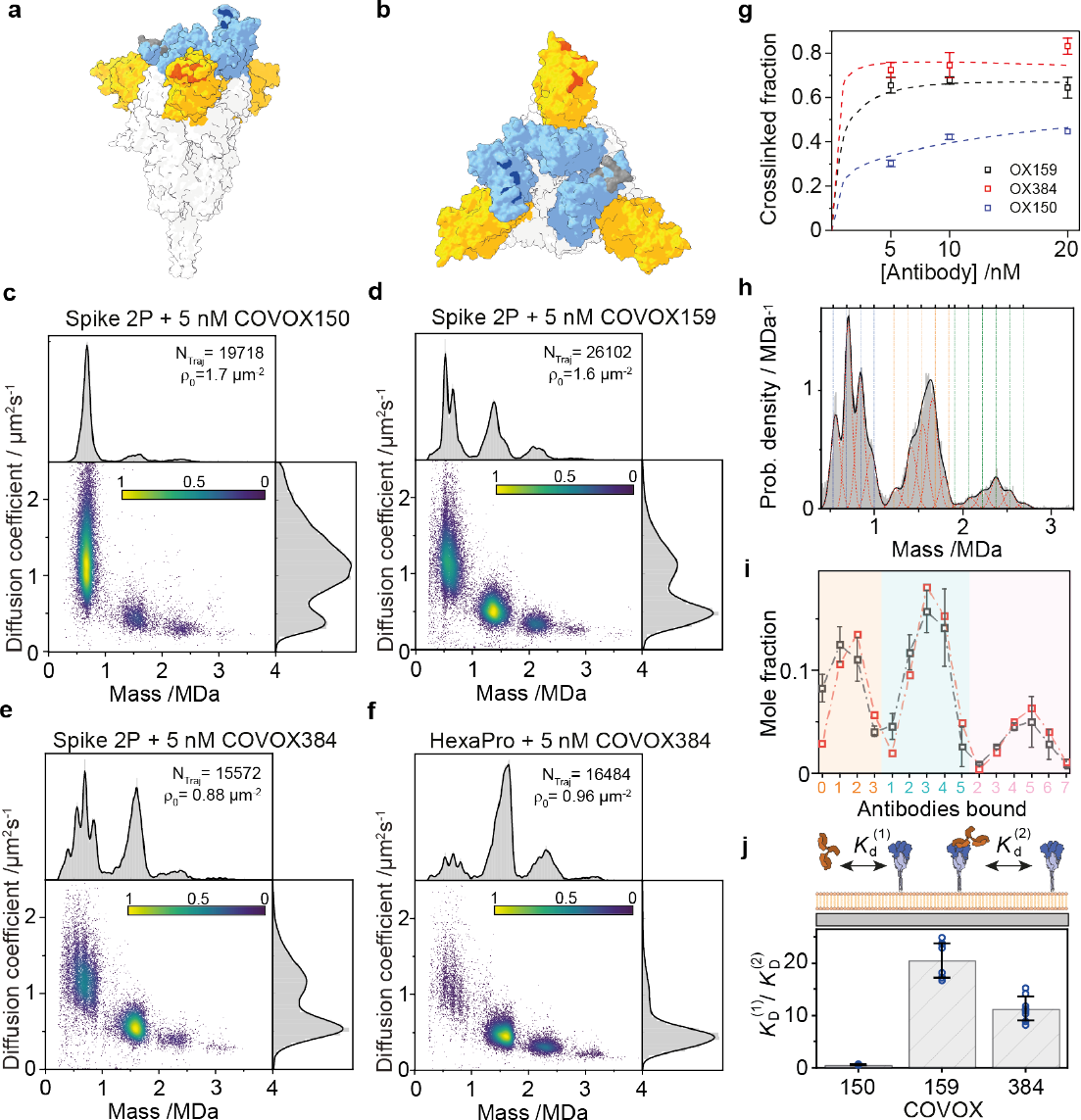
Patient-derived antibodies use induced oligomerisation for both inhibition and enhancement of inhibition. (**a**,**b**) Side and top view of the spike trimer based on PDB entry 6vsb (*2, 3*) with the NTD (yellow) and RBD (light blue) domains highlighted. The binding locations of the different antibodies are highlighted based on their identified interactions (*5*). Grey, dark blue and red correspond to COVOX384 (“down” state of the RBD), COVOX150 (“up” state of the RBD) and COVOX159, respectively. (**c-e**) Two-dimensional plots of diffusion coefficient *vs*. mass for equilibrated tethered wtSpike with added 5 nM of (**c**) COVOX150, (**d**) COVOX159 and (**e**) COVOX384. The number of trajectories sampled for each solution condition and the measured initial surface density of Spike on the supported lipid bilayer are indicated for each plot. (**f**) COVOX384 interacting with tethered wtSpike, stabilised by 6 proline substitutions (HexaPro). (**g**) Mole fraction of spike trimers in higher oligomeric states as a function of the concentration of added antibody to the solution. Values were extracted by counting the number of trajectories for each oligomeric state relative to the total number of detected trajectories while considering the stoichiometry of each state. Symbols and error bars indicate the average values and standard deviation of two independent measurements (except for the case of COVOX150 at 20 nM that includes one repetition). Coloured dashed lines corresponds to our fitted 2D thermodynamic model (**SI Section 4**) (**h**) Representative normalised mass distribution of equilibrated, tethered wtSpike, with 10 nM COVOX159, fitted to a sum of individual Gaussian functions constrained around the expected mass of the different spike:antibody stoichiometries. The black line corresponds to the sum of the Gaussian functions and red dotted lines to the individual Gaussian fits (see **SI Section 4** for complete description of fitting procedure). Dashed vertical lines indicate the masses of the expected wtSpike:antibody complexes, where blue, orange and green correspond to complexes containing one, two or three spike trimers, respectively. (**i**) Mole fractions of individual complexes (grey symbols), calculated from the fitted relative areas of the individual Gaussian functions from **h**. The x-axis corresponds to the number of bound COVOX159 antibodies to an individual spike trimer (orange), to two trimers (cyan) or to three trimers (pink). Error bars indicate the variation between two independent experiments. Red symbols correspond to the predicted mole fractions based on our globally fitted 2D thermodynamic model. (**j**) Ratio of binding affinity from solution, 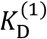, to its two-dimensional affinity for crosslinking, 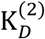. Both values were calculated from the fitted interaction free energies (**SI Section 4**). Average values for the different antibodies are indicated by the hight of each column, where the statistical variation of the fitted ratio values (blue symbols) was calculated by repeating the fitting procedure (global fitting for 5,10, and 20 nM of antibody simultaneously) for different sets of concentrations and repetitions (3 different concentrations of antibody per set and two independent repetitions for each concentration, except for 20 nM COVOX150). Standard deviations are indicated by black error bars.

The ability to identify and localise individual trimers as a function of time also enables us to quantify their mobility. Plotting the cumulative distribution function of observed step sizes yields an accurate estimate of the diffusion coefficient, on a trimer-by-trimer basis through appropriate fitting (**Fig. 2d**). The measurement precision for individual complexes is mainly limited by trajectory length, which is determined by the achievable field of view, the mobility and our ability to link trajectories in the presence of many molecules simultaneously diffusing on the bilayer. The observed mobilities are Brownian in nature, as expected for individual lipids diffusing in an idealised bilayer membrane.

We can then combine the resulting mass and mobility measurements from individual recordings to assess the overall behaviour of the system, where each data point in the two-dimensional scatter plot corresponds to a single trajectory (**Fig. 2e-g**). For wtSpike in the absence of ACE2, we find highly homogeneous behavior dominated by individual trimers with an average mass of 494 kDa diffusing with D = 1.2 ± 0.3 μm^2^/s. The measured mass on the bilayer is slightly lower than that obtained from a regular MP assay (548 kDa), because of the effective height and flexibility of the spike trimer. This leads to variations in the pathlength between reflected and scattered light that forms the basis of the optical contrast in MP, with the consequence of lowering the optical contrast (**SI Fig. 5**). We measured and corrected this variation using measurements on glass and comparison with a calibration protein on the supported lipid bilayer (**SI Fig. 5**), resulting in corrected mass histograms (**Fig. 2e-g**). The mass distribution of species is now dominated by trimers, because the bilayer serves to specifically pull down his-tagged species, and non-covalent oligomers of trimers likely disassemble during the experiment.

**Figure 5.**
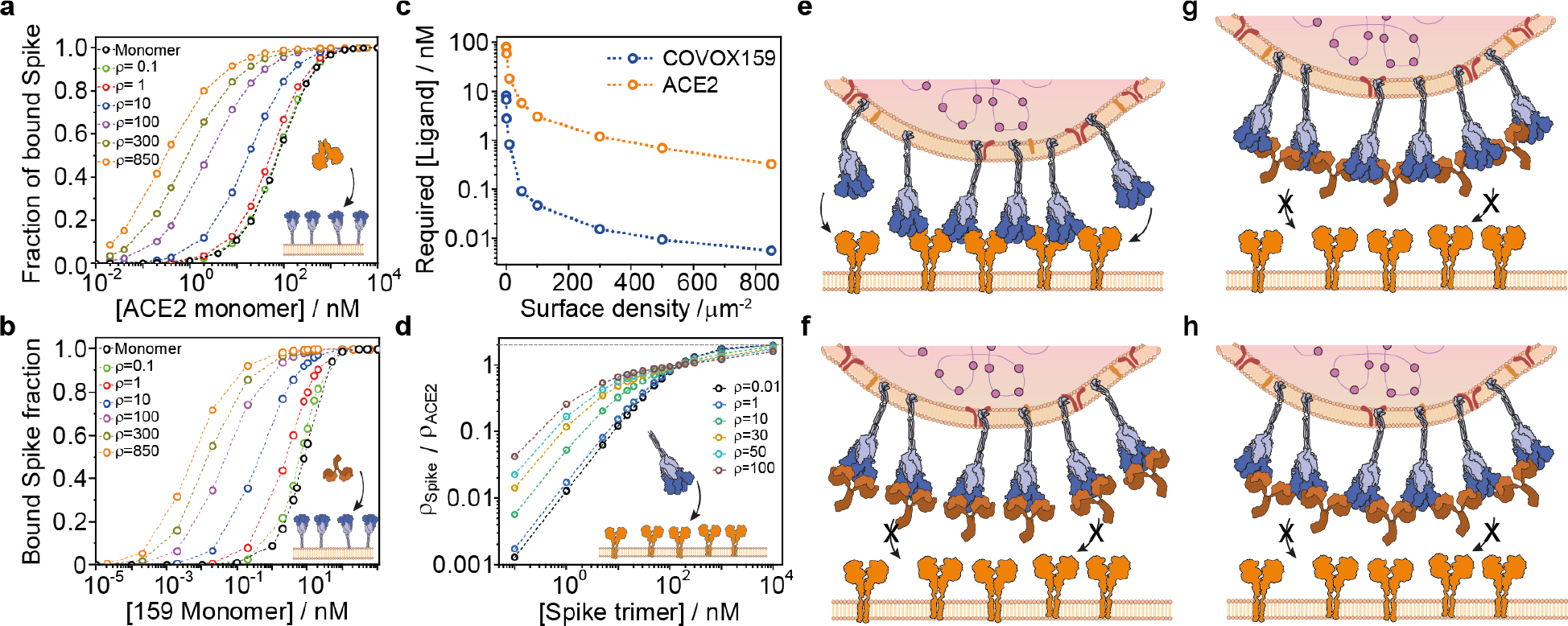
Induced oligomerisation enhances binding of multivalent ligands to their surface multivalent receptors. (**a**,**b**) Molar fraction of Spike trimers bound to at least one ACE2 (**a**) or COVOX159 antibody (**b**) for different spike surface densities calculated from experimentally determined affinities. The monomer binding curve (black), corresponds to no induced oligomerisation. (**c**) Required ligand solution concentration resulting in half of spike bound to at least one ligand as a function of spike surface density. (**d**) Normalised surface density of spike bound to a surface containing different densities of diffusive ACE2 dimers as a function of spike solution concentration. (**e**-**h**) Mechanisms of binding and inhibition of SARS-CoV-2 to its host cell surface. (**e**) Induced oligomerisation of spike and ACE2 during cell-surface binding. (**f**) Inhibition of SARS-CoV-2 binding by blocking the ACE2 binding site by competitive antibodies. (**g**) Blocking cell-surface attachment without affecting the ACE2 binding site by spike oligomerisation alone. (**h**) The most potent antibodies combine ACE2 binding site blocking with spike crosslinking in their mechanism of action.

The addition of 100 nM ACE2 causes clear changes to these distributions. As expected from our solution-based measurements, we now observe wtSpike with 0, 1 and 2 ACE2 bound, but with a clear additional distribution above 1 MDa in the region expected for dimers of spike crosslinked and bound by ACE2, which is also associated with a drop in the diffusion coefficient (**Fig. 2f**). These species appear at a mass in good agreement with the expected mass for the respective spike-ACE2 oligomers, which are evident in both the individual trajectories, and when inspecting the overall envelope of measured diffusion coefficients. Repeating these experiments at similar ACE2 concentration and spike density with the Omicron variant of spike (omSpike) results in much more extensive (and rapid) oligomerisation (SI Movie 3), including signatures consistent with ACE2 decorated spike trimers (**Fig. 2g**). This increase in oligomerization propensity is also evident in the rise of the low mobility shoulder in the overall envelope of measured diffusion coefficients.

The simultaneous measurement of mass and mobility of individual spike complexes in the presence of ACE2 on lipid bilayers provides further evidence for ACE2-induced oligomerisation of spike. Species consistent with a dimer of spike exhibit roughly half the mobility of the monomer, with a further proportional reduction observable for the trimer, and even tetramer. Plotting the diffusion coefficient as a function the reciprocal of the number of spike trimers yields a linear relationship (**Fig. 2h and SI Fig. 6**). Such behaviour is expected for a digital increase in the number of bound lipids per complex, with larger oligomers slowing down due to the increase in drag experienced by multiple lipids incorporated into the bilayer membrane.

### Binding of ACE2 to Spike exhibits variant-dependent cooperativity

Our results demonstrate that ACE2 oligomerizes spike both in solution and on lipid bilayers, and suggest that the ACE2:spike interaction is much weaker than previously reported. At the same time, we have not yet provided a full quantification of the molecular interactions with stoichiometric resolution. To take advantage of this unique capability of MP, we repeated the experiments reported in **Fig. 1**, but mixed ACE2 and spike at nM concentration, which enables us to avoid the formation of large oligomers and loss of spike from solution (**Fig. 1**). For mACE2, we find clear signatures of 1,2 and 3 bound mACE2 to spike, with the abundance of higher occupancy increasing as expected with mACE2 concentration (**Fig. 3a and SI Fig. 7**). Quantifying the fractional occupancy of each binding site enables us to fit the data globally, yielding *K*_D_ = 170 ± 5 nM (see SI for a description of the fitted model). Fitting the data to a model allowing for cooperativity between the 3 binding sites (**Fig. 3b**, solid) does not lead to a significant improvement over a simple binding model, suggesting that the three RBD binding sites are independent for mACE2 binding. The affinity of spike is substantially weaker than that for RBD to mACE2, which is likely a result of additional steric constraints encountered in the full trimer, including the possibility of the RBD to be orientated in a “up” or “down” conformation (*7*).

Titrating ACE2 *vs* spike results in improved peak separation due to the doubling in ligand mass (**Fig. 3c**). As previously, we observe an increase in ACE2 occupancy with increasing concentration. Contrary to our results with mACE2, however, we could not detect significant amounts of spike decorated with three ACE2 (**SI Fig. 8**), indicating substantial steric inhibition of the final binding site in the presence of two bound ACE2. Comparison of the 50 nM ACE2 with 150 nM mACE2 traces shows similar relative amounts of free and single ACE2 bound spike, but much lower amounts of doubly bound ACE2 compared to mACE2, indicative of negative cooperativity affecting binding of the second ACE2 to a singly occupied trimer.

To quantify these effects, we can turn to the same approach previously applied to mACE2, and fit the fractional occupancies as a function of ACE2 concentration to a simple model with the same interaction energy for all binding sites (**Fig. 3d**, dashed), as well as a more complex one allowing for cooperativity (**Fig. 3d**, solid). The difference in the model’s ability to reproduce the data (**Fig. 3d** solid vs dashed) demonstrates the statistical justification for including a cooperative component to the interaction. The global fit to the data yields *K*_D_ = 42 ± 2 nM (84 ± 4 nM in monomer concentration) for the first binding site and negative cooperativity of δΔG°= 0.68 ± 0.04 kcal mol^-1^. Repeating these experiments with omSpike exhibits similar behaviour in terms of a maximum of two ACE2 bound, but binding of the second ACE2 now exhibits positive cooperativity, which can be seen in both the ability to reproduce the measured distributions and the titration data (**Fig. 3e**,**f and SI Fig. 9**). We obtain *K*_D_ = 82 ± 3 nM (164 ± 6 nM in monomer concentration) for the first binding site and positive cooperativity of δΔG°= -0.44 ± 0.04 kcal mol^-1^.

These results quantify the interaction between ACE2 and spike trimers at a molecular level. We find that the 1:1 interaction is roughly two orders of magnitude weaker than previously reported using conventional surface-based assays. At the same time, ACE2 binds more strongly to spike than mACE2, suggesting additional stabilisation for the full dimer. For both spike variants tested, we find significant cooperativity that switches from negative to positive from wtSpike to omSpike. Interestingly, the measured 1:1 molar affinity for omSpike is half as strong as for wtSpike, contrary to previous reports and the hypothesis that tighter interactions correlate with enhanced infectivity.

### Induced oligomerisation dominates the interaction of spike with patient-derived antibodies

Given our observations of cooperativity and oligomerisation for the ACE2-spike interactions, we wondered if they play a role in antibody-spike interactions, given that antibodies provide dual binding potential, in principle similar to ACE2. In addition, we wondered if these effects can help explain and understand differences in behaviours between patient-derived antibodies. We chose three previously studied antibodies(*5*) based on their representative behaviour with respect to infectivity inhibition and receptor binding, differences between Fab fragments and full-length IgG, and the fact that they are reasonably representative of the types of behaviour seen across a large range of antibodies: 1. Exhibiting only small differences between Fab and IgG for neutralisation and binding (COVOX150). 2. Complete lack of neutralisation for Fab (COVOX159). 3. Strongly enhanced neutralisation and binding for the IgG over Fab (COVOX384). COVOX159 further differs from the other antibodies in that its epitope is in the NTD domain (**Fig. 4a**,**b**, red), not in the RBD domain (**Fig. 4a**,**b**, dark blue) and thus does not per se block binding to ACE2.

When adding 5 nM COVOX150 to wtSpike in a bilayer-based assay, almost all spike binds to a single antibody (**Fig. 4c and SI Fig. 10**), indicative of a very strong 1:1 interaction. Nevertheless, the degree of induced oligomerisation is comparatively weak, roughly matching that observed for ACE2 and omSpike, despite the latter exhibiting approximately two orders of magnitude weaker 1:1 affinity. Compared to COVOX150, COVOX 159 induces oligomerisation much more strongly, despite what appears to be a weaker 1:1 affinity, given that we can still observe free spike trimers at the same antibody concentration (**Fig. 4d**). The scatter plot of molecular mass and mobility now contains strong signatures of monomer, dimer, trimer and tetramer of spike, with the dimer most abundant. For the first time, the distribution of mobilities is now dominated by oligomerised spike. Compared to COVOX150, there is a correlation between the abundance of multiple antibodies bound to a single spike trimer, and the degree of oligomerisation. This behaviour is similar for COVOX384 (**Fig. 4e**), which exhibits an almost identical distribution of species and mobilities to COVOX159. Oligomerisation almost completely dominates the interaction when combining COVOX384 with the HexaPro variant of spike(*1*), which maintains the RBD predominantly in the “down” position (SI Movies 4-5, **Fig. 4f**) (*27*).

From these data (**SI Fig. 10-12**), we can directly determine the crosslinked fraction of spike, which shows a clear trend from COVOX150 to COVOX159 and COVOX384 (**Fig. 4g**). We can also determine the amount of free and bound spike-antibody oligomers by fitting a set of Gaussian functions, constrained to the expected masses of the different oligomers (**Fig. 4h, SI Fig. 10-12**), from which we can extract the mole fraction of each species on the bilayer surface (**Fig. 4i**, grey). Based on these mole fractions and their variation with antibody concentration (**SI Fig. 13**) we can use a comprehensive thermodynamic model that takes into consideration oligomerisation on the membrane, its conformational degeneracy (*28, 29*) and the interactions between spike and soluble antibodies (**see SI**) to quantify the energetics, affinities and degree of cooperativity for each antibody-spike system (**Fig. 4i**, red).

Fitting the detected mole fractions to the model allows us to determine both the 3D (soluble antibody to surface bound spike) and the 2D (antibody-bound spike to free spike) standard free energies of interaction (**SI Table 2**). While all antibodies exhibit primary 3D affinities in the 1-5 nM regime (**SI Table 2**), both COVOX159 and COVOX384 oligomerise spike much more strongly than COVOX150, as seen by the ratio between their 3D and 2D *K*_*D*_s (**Fig. 4j**). The observed behaviour is directly linked to the inhibition efficacy of the antibodies. COVOX 150, which oligomerises weakly but binds to the RBD, exhibits the overall weakest IC50 of the three antibodies tested, with small differences between binding, inhibition, Fab and IgG (discussed further below). COVOX384 binds to the RBD and oligomerises strongly, making it one of the most potent antibodies (*5*). Remarkably, COVOX159 does not bind to the RBD region, and thus does not inhibit ACE2 binding by occupying the RBD, yet exhibits strong inhibition, which is likely entirely attributable to its ability to oligomerise spike on the surface of the virus. In fact, ACE2 still binds successfully to a COVOX159-prebound spike in solution (**SI Fig. 14**)

Our approach is limited to spike densities on the membrane on the order of 1-2 μm^-2^ due to the need to separately visualise and quantify individual spike complexes in a diffraction-limited imaging system. The density on the surface of the virus, however, is about three orders of magnitude higher than the experimental limit in this study, with structural flexibility that may further facilitate multivalent interactions (*30, 31*). Having quantified the relevant energetics at the molecular level, we can now use the thermodynamic parameters, in this case taken from the spike-COVOX159 or wtSpike-ACE2 systems, to deduce macroscopic observables such as the fraction of bound RBD and the fraction of bound spike as a function of ligand concentration, multivalency and different spike surface densities.

The thermodynamic model, predicts a >2 orders of magnitude improvement in RBD occupancy (**SI Fig. 15**) or bound spike fraction (**Fig. 5a**,**b**) for an oligomerisation-prone binder compared to the monomer (black curves) when surface densities approach the density of spike proteins on the virus surface for both ACE2 and COVOX159 (**Fig. 5a**,**b**). This effect is strongest at low ligand concentrations (< *K*_*D*1:1_), where oligomerisation dominates and improves spike occupancy even at extremely low ligand concentrations (**Fig. 5c**). At higher ligand concentrations the oligomerisation capacity is already saturated and limited by free spike surface density, therefore the binding becomes dominated by the 3D affinity, which leads to the observed convergence of binding curves (**Fig. S15**).

Finally, we show that cooperative oligomerisation enhances spike binding to a membrane surface containing different densities of ACE2 (**Fig. 5d**). Here, we assumed the interaction parameters of omSpike, where each spike trimer can bind up to two ACE2, and the second binding event includes positive cooperativity. In this case spike and ACE2 can form linear oligomers on the membrane surface. Similar to previous results and depending on ACE2 surface density, oligomerisation results in more than one order of magnitude enhancement of spike affinity. The enhancement is maximal at a concentration much lower than the fundamental *K*_*D*_ (82 nM), while at higher concentrations than the fundamental *K*_*D*_ the trend is inverted owing to the larger number of binding sites for un-oligomerised ACE2. In all cases, our calculations reveal the role of cooperativity and multivalency in strengthening the apparent affinity up to several orders of magnitude compared to the fundamental *K*_*D*_ of the monomeric subunits. The increase in the binding free energy gain resulting from induced oligomerisation is a function of the 2D affinity (oligomerisation propensity) and the chemical potential of the free tethered component and therefore its surface density. Thus, we expect the binding enhancement to be most prominent at high surface densities and solutions concentrations lower than the solution *K*_*D*_, while at high concentration this enhancement factor is lower since the ligand is already saturating the tethered receptor owing to its 3D affinity.

## Discussion

Our results have significant implications for our quantitative molecular understanding of spike-ACE2 and spike-antibody interactions specifically, but also for oligomeric protein-protein interactions more broadly. The ability of MP to disentangle protein-protein interactions and avidity on a subunit-by-subunit level reveals a much more nuanced picture compared to that emerging from bulk biophysical and structural methods. Our observed ACE2-RBD affinity of 28 ± 9 nM fits well into the range of reported values (17 -75 nM) (*7, 8, 11, 12, 22-24*). However, we find 3- and 6-times weaker binding to spike for both variants tested contrary to previous reports, which reported up to 100-fold enhancement of the interaction strength (*11*), down to apparent *K*_*D*_s of 0.015 – 3 nM (*11, 13, 25, 26*) when both ACE2 and spike are multivalent. A lower interaction strength, however, is expected for spike in the absence of intramolecular avidity, given that the RBD can occupy an “up” or “down” position, only the latter of which is capable of binding ACE2. A likely explanation for this difference, and for many of the discrepancies between our observed affinities and those reported with bulk methods, can be attributed to the fact that bulk characterisation can inadvertently allow for intermolecular interactions that have not been taken into account. Thus, our results agree well with assays using monomeric constructs, but deviate substantially with those where oligomeric ligands are added to a surface covered with receptors.

We show that the multivalency of both spike and ACE2 plays an important role in their interaction. Our observations of ACE2-induced oligomerisation of spike both in solution and on lipid bilayers provide insight into its importance for both cellular binding and internalisation, as well as inhibition. Previous reports have demonstrated two orders of magnitude improvement in efficacy of dimeric over monomeric ACE2 in the context of viral inhibition (*11, 16, 17*). This, could *a priori* be explained by intra-spike avidity were a dimeric ACE2 can bind to two RBD on the same spike, resulting in enhanced affinity. Yet, despite the anticipated high stability of such a complex, the associated structure has never been reported. Additionally, intra-spike avidity would block extended spike oligomerisation, because it requires at least two inter-spike interactions per trimer. Our results allow us to rationalise the molecular mechanism underlying the inhibitory efficacy of multivalent ligands, while being in quantitative agreement with previously reported enhancement in inhibition. Furthermore, the ability of ACE2 to oligomerise spike makes it essentially impossible for the virus to detach from the cellular surface due to the associated avidity effects and drives receptor clustering, which is associated with signalling for internalisation and inducing membrane curvature and ultimately fusion (*32, 33*).

In contrast to standard 1:1 affinities, we resolve the stoichiometries of the interaction and their relative abundances, which provide insight into cooperative effects. We find binding to be independent for mACE2 and wtSpike, but substantial effects for ACE2 that changes with the spike variant. Contrary to previous reports (*11, 13*), we find that the 1:1 affinity of the Omicron variant is weaker than that of the original Wuhan-Hu-1 strain, but exhibits a switch from negative (Wuhan) to positive (Omicron) cooperativity for higher binding stoichiometries, leading to increased oligomerisation propensity. This combination will lead to preferential binding to tissues and cells with higher ACE2 density, where multiple interactions with ACE2 are favoured (**Fig. 5d**,**e**).

Our results with patient-derived antibodies reveal a hierarchy of interactions and help explain mechanisms of virus neutralisation. COVOX150 exhibits strong binding to the RBD but weak oligomerisation (**Fig. 5f**), leading to relatively weak inhibition (IC50 = 0.15 nM) (*5*). COVOX159 exhibits slightly better inhibition (IC50 = 0.033 nM) (*5*) despite its slightly weaker affinity and leaving the RBD untouched, which likely arises from its ability to efficiently induce the oligomerisation of spike trimers on the viral surface. The best performing antibody (COVOX384) combines RBD binding and oligomerisation for overall maximal inhibition (IC50 = 0.013 nM) (*5*), despite similar 1:1 binding affinity to the other antibodies. The dramatic improvements in target occupancy achievable at low ligand concentrations (**Fig. 5a**,**b**), the ability to achieve an inhibitory effect without targeting the RBD, and the ability to enhance the effect of binding to the RBD (coupled with the intrinsically dimeric structure of antibodies) raises the possibility that induced oligomerisation is a more general inhibitory mechanism used by the immune system.

Taken together, our results reveal the nature and importance of induced oligomerisation for both virus-cell binding and inhibition by antibodies. They demonstrate that models solely based on 1:1 interactions fail to identify to the molecular details that are responsible for fundamental differences in infectivity and inhibition. Instead, propensity to induce oligomerisation is as important if not more important than the strength of the molecular interfaces defining the fundamental interaction. Our results also raise the question to which degree oligomerisation represents a more fundamental mechanism driving interactions and effects such as tropism. In addition, our results raise the possibility that these interactions are in fact fundamental to the way antibodies function, explaining their unique, effectively dimeric structure. Importantly, these effects do not rely on multivalency alone, they require geometric restrictions, which in this case prevent both ACE2 and antibodies to bind two sites within the oligomeric target and instead drive oligomerisation, an effect that could be leveraged in the rational design of future therapeutics.

## Acknowledgments

The pCAGGS expression plasmid encoding the Wuhan-hu-1 2P spike trimer and the vector encoding the Wuhan-hu-1 spike receptor binding domain (RBD) were kindly provided by the Krammer Laboratory, Department of Microbiology, Icahn School of Medicine at Mount Sinai, New York. The pHL-sec vector encoding Omicron 2P spike was a kind gift from the Townsend Laboratory, Weatherall Institute of Molecular Medicine, University of Oxford. pcDNA3-sACE2-WT(732)-IgG1 was a gift from Erik Procko (Addgene plasmid #154104; http://n2t.net/addgene:154104; RRID:Addgene_154104).

## Funding

European Research Council PHOTOMASS 819593 (PK)

Engineering and Physical Sciences Research Council EP/T03419X/1 (PK) UKRI Future Leaders Fellowship MR/V02213X/1 (WS)

University of Oxford’s COVID-19 Research Response Fund (WS & PK) EMBO Postdoctoral Fellowship ALTF 198-2020 (RA)

Oak Foundation (JM, GRS)

Wellcome trust SEACOVARIANTS consortium 226120/Z/22/Z (JM, DS, GRS) Wellcome Trust Centre for Human Genetics 090532/Z/09/Z (JM, GRS)

The Chinese Academy of Medical Sciences Innovation Fund for Medical Science, China 2018-I2M-2-002 (LC, JM, GRS)

Fast Grants, Mercatus Center, NIH Research Biomedical Research Centre Funding Scheme (GRS)

Medical Research Council (MR/N00065X/1) (DS)

Schmidt Futures, and Red Avenue Foundation (GRS)

Wellcome Trust PhD Programme 203853/Z/16/Z (JB)

Oxford Glycobiology Endowment (NZ)

Biotechnology and Biological Sciences Research Council BB/V011456/1 (JLPB, WBS)

## Author contributions

Conceptualization: RA, AO, JLPB, WBS, PK

Methodology: RA, AO, WBS, PK

Investigation: RA, AO

Visualization: RA, WBS, PK

Computational support: LPR, MD

Supply of antibodies: LC, WD, PS, JM, DZ, DS, GRS

Supply ACE2 and Spike constructs: SV, JB, MH, MH, NZ

Funding acquisition: RA, PK

Project administration: PK Supervision: PK

Writing – original draft: RA, PK

Writing – review & editing: RA, WBS, JLPB, PK

## Data and materials availability

All data necessary to support the conclusions are available in the manuscript or supplementary materials and are deposited in the University of Oxford Research Archive.

### Open access

This research was funded in whole or in part by the ERC (PHOTOMASS 819593) EPSRC EP/T03419X/1. For the purpose of Open Access, the author has applied a CC BY public copyright license to any Author Accepted Manuscript (AAM) version arising from this submission.

