## Supplementary Information for "Cooperativity and induced oligomerisation control the interaction of SARS-CoV-2 with its cellular receptor and patient-derived antibodies"


**The PDF file includes:**

Materials and Methods
Figs. S1 to S21
Tables S1 to S2
References

**Other Supplementary Materials for this manuscript include the following:**

Movies S1 to S5

### Materials and Methods

Buffers lipids and commercial proteins

HEPES (H3375), KCl(P9541) and Chloroform (288306), Magnesium Chloride hexahydrate (M2670), Sodium Chloride (S3014) and Trizma base (T1503) were purchased from Merck Life Science UK Limited. Dulbecco's Phosphate Buffered Saline (DPBS) was purchased from ThermoFisher Scientific. UltraPure 10% SDS (15553-035, Invitrogen). Lipids, 1,2-dioleoyl-snglycero-3-phosphocholine, 850375P (DOPC); 1,2-dioleoyl-sn-glycero-3-[(N-(5-amino-1-carboxypentyl)iminodiacetic acid)succinyl] (nickel salt), 790404P (DGS-NTA) and 1,2-dioleoyl-sn-glycero-3-phosphoethanolamine-N-[methoxy(polyethylene glycol)-2000](ammonium salt), 880130P (DOPE-PEG2k) were purchased from Avanti Polar Lipids. The lipid powders were kept at -80 °C and aliquots were stored in chloroform at −20 °C. Proline stabilised Wuhan-Hu 1 HexaPro spike was purchased from AcroBiosystems (Cat.# SPN-C52H9).

Protein expression constructs

The pCAGGS expression plasmid encoding the Wuhan-Hu-1 2P spike trimer was a kind gift from the Krammer Laboratory, Department of Microbiology, Icahn School of Medicine at Mount Sinai, New York (*1*). The vector encoding the Wuhan-Hu-1 spike receptor binding domain (RBD) was a kind gift from the Krammer Laboratory, Department of Microbiology, Icahn School of Medicine at Mount Sinai, New York (NR-52309). The pHL-sec vector encoding Omicron 2P spike was a kind gift from the Townsend Laboratory, Weatherall Institute of Molecular Medicine, University of Oxford. Expression constructs encoding monomeric (a.a. 19-611) and dimeric (a.a. 19-726) ACE2 were made using gene the pHL-sec vector and the ORF cDNA cloning vector from Stratech Scientific (HG10108-M-SIB). pcDNA3-sACE2-WT(732)-IgG1 was a gift from Erik Procko (Addgene plasmid #154104; http://n2t.net/addgene:154104; RRID:Addgene_154104)(*2*).

Protein expression and purification

Proteins were transiently expressed in HEK293F (FreeStyle™, Thermo Fisher Scientific) cells. Cells were cultured in Freestyle 293 expression media (ThermoFisher Scientific) and incubated at 37 °C, 8% CO2 and 120 rpm. Transfection was achieved using FreeStyle™ MAX reagent (Invitrogen) and OptiMEM™ (Gibco) following a published protocol (*3*). Five days post transfection, cell culture supernatant was harvested by centrifugation at 3000 x g for 10 min and then filtered using 0.45 µM pore size filters (Merck). Supernatants were supplemented with 10 mM imidazole and His-tagged proteins were purified using a HisTrap HP, 5 mL column (Cytiva) connected to an ÄKTA pure protein purification system (Cytiva). ACE2 containing IgG1-Fc was purified using a HiTrap Protein A HP, 1 mL column. Proteins were further purified by size exclusion chromatography (SEC) using a Superose 6 increase 10/300 GL column (GE Healthcare) equilibrated in Dulbecco's phosphate-buffered saline (DPBS, pH 7.4, ThermoFisher Scientific). Protein containing SEC fractions were pooled and concentrated using Amicon molecular weight cut-off centrifugal filters (GE Healthcare). Protein concentrations were determined using a Nanodrop spectrophotometer (Thermo Fisher Scientific) at absorbance 280 nM and corrected for protein molecular weight and extinction coefficient.

Solution mass photometry measurements

The interaction between wtSpike and ACE2 at $\mu$M concentration as well as the interaction between soluble RBD and ACE2 were conducted using a OneMP (Refeyn Ltd., Oxford) with a field of view of 3.8×10.7 $\mu m^2$. Measurements were conducted using microscope glass coverslips

(24 × 50 mm, Menzel Gläser, VWR 630-2603) that were cleaned by three consecutive cycles of 5 min bath sonication in Milli-Q® water (18.2 MΩ·cm) (Milli-Q), 50% isopropanol in Milli-Q, and Milli-Q again. Cleaned coverslips were then dried using nitrogen flow and 3 mm silicone gaskets (GBL103250, Grace Bio-Labs) were attached on the coverslip surface. The attached gasket was prefilled with 5 µL of buffer and the position of the focus was adjusted for optimal contrast. Measurements were performed using a field of view size of 3.2 by 9 µm, and the frame rate was 1000 Hz followed by frame binning of 5 that resulted with an effective frame rate of 200 Hz.

Measuring the interaction of spike and ACE2 at µM concentration

For spike–ACE2 interaction measurements, wtSpike at a trimer concentration of 0.55 µM was mixed with increasing concentrations of soluble ACE2 (a.a 19-726), from 0 to 3.3 µM or soluble monomeric ACE2 (a.a 19-611) at concentration of 1.65 µM. The mixed proteins as well as the individual components were incubated at ambient room temperature for 10 min. Following incubation, the solutions were rapidly diluted in DPBS buffer to the measured concentrations of 16.7 nM wtSpike trimer and between 0 and 100 nM ACE2. Then, 30 µL of the diluted sample was loaded to the measurement gasket. Data acquisition was started within 14–18 sec following dilution in order to minimise complex disassociation. Data was recorded for 60 sec, and each experiment was repeated between 3 and 6 times (as indicated in **Fig. 1**).

Measuring the interaction between soluble RBD and ACE2

The ACE2 - RBD interaction was measured by mixing in DPBS 10 nM of ACE2 (a.a 19-726) with increasing concentration of between 0 and 30 nM of soluble RBD (**Fig. S2**). The mixture was incubated at ambient room temperature for 10 min, then the sample was added to the gasket and data was recorded for 60 sec. Data presented in **Fig. S2** represents between 2 and 4 technical repeats for each mixing condition.

Solution mass photometry measurements of Spike-ACE2 and Spike-antibody interaction at nM concentration

MP experiments at nM concentration (wtSpike-mACE2/ACE2 and omSpike-ACE2 interactions) were carried out on a commercial mass photometer (TwoMP, Refeyn Ltd.). Spike, at trimer concentration of 25 nM, was incubated with ACE2 at its indicated concentrations and at room temperature for between 5 and 40 min prior to each measurement (**Fig. S7-9**). For each solution condition, between 3-5 technical repeats were taken at consecutive times, which show no time variation in the resulting histograms. All samples were measured in DPBS. For all measurements, the coverslip, attached to a gasket, prefilled with 10 µL of buffer was placed on top of the microscope stage and the focus position was adjusted. Immediately prior to each acquisition, 10 µL of protein solution was added to 10 µL of buffer in a single gasket. A 60 sec video was immediately recorded following the addition of the protein sample, capturing landing events of individual proteins on the coverslip surface. The timescale from loading to the beginning of the acquisition was <10 s. Examination of the distribution of oligomers during the 60 sec data acquisition showed no significant variation in the normalized distribution of the interacting species suggesting that any possible disassembly process that may take place following the addition of protein to the gasket is slower than our measurement timescale and thus, the measured distribution corresponds to the equilibrated solution distribution. For all spike-ACE2 measurements a standard field of view was used ($10.9 \times 4.3\ \mu m^2$) with an acquisition frame rate of 500 Hz, which followed by frame binning of 2, resulting in an effective frame rate of 250 Hz. An identical procedure was followed to measure the co-binding of antibody COVOX159 and ACE2 to wtSpike, with the exception that the large field of view (16.9 × 12.0 μm$^2$) was used, which allows for improved statistics owing to the larger surface area of detection. This resulted in a frame rate of 130 Hz.

Solution mass photometry data analysis

Data analysis was performed with DiscoverMP v2023R1.2 (Refeyn Ltd.) in which rolling ratiometric movies were generated using an averaging window size of 5 frames (20 and 38 ms integration time for the regular and large FOV respectively). Threshold parameters for particle detection were set to their default values of 1.5 (threshold 1) and 0.25 (threshold 2). For each data set, the calibration of interferometric contrast to mass was performed using a protein standard while using the same acquisition parameters, similar to the previously reported procedure (*4*).

Distribution analysis of solution mass measurement

Mass measurement of individual particles performed by DiscoverMP were further analysed to extract interaction parameters using custom-written python scripts.

RBD-ACE2 interaction landscape

A probability density was generated using a kernel density estimator (KDE) from mass histograms of each technical repeat using the Scikit-learn python package (*5*), with a bandwidth of 3 kDa. Individual KDEs were then averaged to obtain the average normalised probability density  and its standard deviation as a function of mass. A set of six averaged KDEs was than generated for the six ratios between ACE2 and RBD. The model for the interacting subunits was based on four equilibrium reactions describing the possible interactions as is illustrated in **Fig. S2a** including the following,

$$mACE2 \; + \; mACE2 \rightleftharpoons \; \mathrm{ACE2}$$

$$mACE2 \; + \; RBD \rightleftharpoons \; \mathrm{m}ACE2 \; \cdot RBD$$

$$ACE2 + RBD \rightleftharpoons ACE2 \; \cdot RBD_1$$

$$ACE2 \cdot RBD_1 + RBD \rightleftharpoons ACE2 \cdot RBD_2 \; .$$

Since all reactions are bimolecular, all dissociation constants are described by the following equation,

$$K_{\mathrm{D}} = \frac{[A][B]}{[AB]} = C_{\circ} \frac{\Omega_{\mathrm{A}}\Omega_{B}}{\Omega_{AB}} e^{\left(-\frac{\Delta G^{\circ}}{k_{\mathrm{B}}T}\right)}$$

Where $\Delta G^{\circ}$ is the standard free energy for the interaction, $C_{\circ}$ is the standard concentration and $\Omega_{\mathrm{i}}$ is the number of equivalent microstates. Here, $C_{\circ}$ was set to 55.5M, and $\Omega$ equals 1 for all species except for the complex $ACE2 \cdot RBD_1$ where its degeneracy is 2.

We can then write the concentration of each in terms of the two basic subunits which are $mACE2$ and $RBD$. We can than write two conservations laws for the molar fractions ($\chi_i = C_i/C_{\circ}$) for the two subunits,

$$\chi_{\mathrm{m}ACE2\,,0} = \chi_{\mathrm{m}ACE2} + \chi_{\mathrm{m}ACE2}^{2} \cdot e^{-\frac{\Delta G_1^{\circ}}{k_{\mathrm{B}}T}} + \chi_{\mathrm{m}ACE2} \cdot \chi_{\mathrm{RBD}} \cdot e^{-\frac{\Delta G_2{}^{\circ}}{k_{\mathrm{B}}T}} + {\chi^2}_{\mathrm{m}ACE2} \cdot \chi_{\mathrm{RBD}} \cdot 2 \cdot e^{-\frac{\Delta G_2{}^{\circ} + \Delta G_1{}^{\circ}}{k_{\mathrm{B}}T}} + {\chi^2}_{\mathrm{m}ACE2} \cdot \chi_{\mathrm{RBD}}^{2} \cdot e^{-\frac{2\Delta G_2{}^{\circ} + \Delta G_1{}^{\circ}}{k_{\mathrm{B}}T}}$$

$$\chi_{\text{RBD},0} = \chi_{\text{RBD}} + \chi_{\text{m}ACE2} \cdot \chi_{\text{RBD}} \cdot e^{-\frac{\Delta G_2{}^\circ}{k_\text{B}T}} + \chi^2{}_{\text{m}ACE2} \cdot \chi_{\text{RBD}} \cdot 2 \cdot e^{-\frac{\Delta G_2{}^\circ + \Delta G_1{}^\circ}{k_\text{B}T}} + \chi^2{}_{\text{m}ACE2} \cdot \chi^2_{\text{RBD}} \cdot e^{-\frac{2\Delta G_2{}^\circ + \Delta G_1{}^\circ}{k_\text{B}T}} .$$

To extract the interaction standard free energy changes, $\Delta G^\circ_{1/2}$, a global fitting procedure was applied to the entire titration data set. Global fitting was done by introducing a normalized Gaussian function (with its area equal to 1), for each of the possible species at its expected mass, while its width was set by our previously measured sample of ACE2 and was set to 10 kDa. The objective function to minimise with respect to the model was the global reduced $\chi^2$ function equal to:

$$\chi^2_{Global} = \frac{1}{N}\sum_{i=1}^{N} \chi^2_{r,i}$$

where, $\chi^2_{r,i}$ is the reduced $\chi^2$ function for the $i$-th solution condition, where the number of free fitting parameters are the two interaction free energies and $N = 6$. At each iteration the model assumed the value of the two free energies and then solved the relative molar fractions of the different species according to the conservation laws. Following this step, the relative concentration of each species at the different mixing ratios was calculated and defined the amplitude of the normal Gaussian function. For each solution condition, a model of the normalized distribution was generated and the model was compared to the relevant KDE while allowing small variations in the centre of mass of each Gaussian to account for possible variations in the focus position and sample drift. The maximum allowed variation in mass was $\pm 10$ kDa and in sigma $\pm 2$ kDa. The results of the fits are shown in **Fig. S2a**. Confidence levels were calculated according to the curvature of $\chi^2_{Global}$ close to its minimal value by assuming that the possible error in $\chi^2_{Global}$ is given by $\frac{\sigma_{\chi^2_r}}{\sqrt{N}}$ (**Fig. S2b,c**). Optimisation was performed using the *scipy.minimize* python package.

<u>wtSpike and omSpike – ACE2 interactions.</u>

As the detected peaks are more resolved owing to the larger mass of interacting species, we can fit individual Gaussian functions directly to the histogram rather than constraining the fitting with the thermodynamic model. We use the same procedure to fit the three data sets for the wtSpike-mACE2, wtSpike-ACE2 and omSpike-ACE2 interactions. For each technical replicate, a KDE was generated for the mass histogram using a bandwidth of size 5 kDa. The KDEs were then fitted to a set of Gaussian functions centered and bound around the expected position of each species, while allowing ±(20-30) kDa variation in the mass during the fit. Here, the intrinsic peak width of the spike trimer peak is larger (~25 kDa), therefore we allowed the width of the Gaussian to vary between 15 and 30 kDa. For all data sets *scipy.optimize.curve_fit* was used for fitting. Only peaks that include spike and its interacting species were fitted. For the interaction involving monomeric ACE2, we set the number of possible species to 4 (free spike, spike bound to 1, 2 and 3 mACE2) as shown in **Fig. S7**. Since spike exhibits small population of dimers of trimers whose mass overlaps with the expected mass of spike-ACE2 complexes, we included more possible species for the interaction between ACE2 and wtSpike including: spike, spike bound by 1, 2 and 3 ACE2 and dimer of spike. **Fig. S8** shows the variation of the fitted peak as a function of ACE2 concentration. Specifically, we enlarge the high mass range, where we indicate the expected position of a dimer of spike (before adding ACE2) that is overlapping with the expected position of spike bound to

two ACE2s. As the ACE2 concentration increases, the overlapping peak shifts towards the expected mass of spike + 2 ACE2. In a similar manner, the shifts in the overlapping peaks of a dimer of spike bound to ACE2 and spike bound to three ACE2 may indicate a very low concentration of spike bound to three ACE2 dimers, however, given the low number of counts compared with the baseline counts its significance is low. The same procedure was followed for the ACE2-omSpike interaction. Here we found a minor amount of impurity in our ACE2 sample that likely results from aggregated ACE2. This fraction did not change with increasing the ACE2 concentration, suggesting it is not concentration dependent aggregation. The impurity appears at 520 kDa close to the mass of unbound spike. While the number of counts is negligible to affect the probability distribution in the low ACE2 concentration measurements (1.5% and 3% for 20 and 50 nM) it could slightly affect the amount of free spike at 70 and 100 nM by a factor of 10% and 20%. We therefore measured the ratio of this peak at different ACE2 concentrations (**Fig. S9**) and corrected the number of counts of the free omSpike according to the counts of the ACE2 multiplied by this ratio. For omSpike we did not find significant evidence for the possibility that 3 ACE2s can bind the same spike trimer, therefore we assume that for Omicron, a maximum of two ACE2s can be bound simultaneously.
From the fitted peaks areas, we could extract the relative occupancy probabilities of the different RBD sites. These probabilities can be fitted to a thermodynamic model taking into account the total concentrations of spike and ACE2 and their interaction free energies similarly to the model used for ACE2-RBD. An illustration of the considered reactions and thermodynamic parameters we used to fit the occupancy probabilities (**Fig. 3d-f**) is shown in **Fig. S17**. The only change between mACE2 and ACE2 binding is the factor of $2^n$ for the number of microstates.

Supported lipid bilayer preparation

Supported lipid bilayers were prepared using a similar procedure as previously reported(*4*). For supported lipid bilayer formation on glass coverslips, phospholipids stocks in chloroform were mixed to form a 10-fold concentrated stock solution with a molar composition of 0.05 mM DGS-NTA, 0.07 mM DOPE-PEG2k and 4.88 mM DOPC (molar ratio of 1/1.4/97.6, respectively). The stock solution was kept at -20 °C. Before use, 50 µL of the lipid stock solution was added to 200 µL of chloroform in a clean glass tube (cleaned by washing with MQ water, followed by 10% SDS and finally with MQ water and acetone before drying). The chloroform was than evaporated by manually rotating the tube while applying a weak flow of nitrogen, followed by 1 h evaporation under vacuum. Following chloroform evaporation, 0.5 mL of buffer (20 mM HEPES pH 7.4, 100 mM KCl) were added to the tube, followed by 2 cycles of 20 min incubation in a 40 °C water bath with vortexing in between. The sealed tube was left at ambient room temperature overnight. The hydrated lipids were then transferred into a 1.5 mL Eppendorf tube and the lipids were tip sonicated using a 2 mm tip probe at 30% power and 1 sec pulse duration separated by 3 sec waiting time for a total of 10 min sonication time (Vibra-Cell, Sonics & Materials). During sonication, the tube was kept on ice. The sonicated lipids were than centrifuged at 21,130 g for 30 min at 4 °C, before removing 0.4 mL of the supernatant.

Glass coverslips were cleaned as described for standard MP measurements and treated with oxygen plasma for 3 minutes at 40% power and 0.6 mbar oxygen pressure (Zepto plasma cleaner, Diener Electronic). Immediately following plasma cleaning, a silicon gasket (GBL103280, Grace Bio-Labs) was placed at the center of the coverslip and 30 $\mu$L of buffer (20 mM Tris pH 7.8, 150 mM NaCl, 2 mM $MgCl_2$) followed by 20 $\mu$L of lipids were added and thoroughly mixed in the gasket. Following 20 min of incubation at room temperature, excess liposomes were washed from solution by multiple washes with DPBS.

Dynamic MP measurements

Dynamic MP measurements of soluble ACE2 (WT(732)-IgG1) or antibodies with tethered wtSpike or omSpike were performed on a commercial mass photometer (oneMP, Refeyn Ltd.). Acquisition used the medium field of view ($6.3 \times 9.9\ \mu m^2$) and at the maximum frame rate of 540 Hz and a metapixel pixel size of 77.35 nm after 4×4 binning. Before each measurement, the washed bilayer was checked for strong scatterers, which are usually immobile/adsorbed vesicles or defects resulting from incomplete fusion on the glass surface. To the clean bilayer, 2 nM of spike trimers were added and the number of diffusing particles were monitored over time to reach a density of about 1 particle per $\mu m^2$. When the desired density was reached, unbound spike protein was washed out by replacing the volume at least 5 times with DPBS. For each tethered spike type (wt, Omicron, HexaPro) we tested its tendency to oligomerize on long timescales without the addition of ligands. In all cases spike did not form oligomers even following several hours of incubation on the supported lipid bilayers. After washing excess spike, either ACE2 or antibodies were added to the solution above the bilayer to reach the desired concentration at a total of 100 $\mu$L volume. A 60 sec measurement was acquired just following the addition of the ligand to record the initial time point of the reaction. For each solution condition, we incubated the bilayer with the proteins for about 1 h, while recording 60 second movies every 5-10 min. Between the measurement, the microscope lid was left open to avoid continuous illumination of the sample, and the sample was covered by a lid containing wet paper towel on its edges to reduce evaporation. When steady state was reached after approximately 40 min, between 3 to 4 pairs of 60 sec measurements were taken, each pair on different areas of the supported lipid bilayer, and 2 min time separation between measurements within the same pair. Before each acquisition, the microscope stage was adjusted to the optimal focus position. Before each data set acquisition, a protein standard was measured to calibrate the contrast to mass conversion using the same acquisition parameters.

Dynamic MP image analysis

All movies were analysed using a custom-written python script based on the previously published package with modifications (*6*). Here, we outline a short explanation of the important steps and input parameters required to analyse dynamic MP data. ***Image processing***: For each analysed movie, all pixels values were first converted to the equivalent photoelectron counts and then each frame was normalized by its total numbers of photoelectrons. The movie was then binned 2-fold to an effective frame rate of 270 Hz, as this reduction sped up analysis time without affecting the contrast sensitivity or resolution given the observed mobilities. Background subtraction of the underlying glass roughness was achieved with moving median ratiometric imaging (*6*), where the difference between each pixel value at time $t$ and a median value calculated for that pixel within a time window of width $T$ centered around $t$ was generated and then divided by the median value. This procedure, allows us to subtract any static background such as glass roughness, while maintaining the dynamic signals originating from diffusing particles. This results in images similar to **Fig. 2b** and **Movies S2-5**. Here, since oligomerized particle diffuse more slowly, we chose a relatively long median window of 600 frames (300 from each side) which is equivalent to 2.2 sec at the given effective frame rate. Following the subtraction of the background a spatial median kernel of size 15×15 pixels was convoluted with each frame to remove any low frequency intensity modulation that originates from rapid scanning of the imaged FOV by the laser. ***Particle detection:*** For particle detection, each image was convoluted with a Laplacian of Gaussian kernel with a sigma of 1.5 pixels. Following the convolution, a threshold value was applied to generate a binary map for candidate pixels exceeding this threshold. Here, we used a threshold value of 0.0015. A second filter was applied to the image that identifies pixels whose value is a local maximum. A candidate pixel for the center of a detected particle is a pixel that fulfilled both criteria. ***Particle fitting:*** Each candidate pixel then serves as the center of a region of interest (ROI) of size 13×13 pixels and the initial guess for the fitting procedure. The contrast of the detected particle is extracted by fitting a point spread function (psf) model to the ROI, by minimizing the square difference between the ROI and the model. The free parameters of the psf model are the 2-dimentional coordinates of its center and its amplitude/maximum contrast. For complete description of the psf model used in this study, see Materials and Methods described in (*6*).

Generating a trajectory from consecutive localisations

To connect individual successful and consecutive fitting events across adjacent frames into individual molecular trajectories of diffusing proteins, similar to the example shown in **Fig. 2c**, we used the linking function of the trackpy python package (*trackpy.link_df*). For our use, there are three relevant parameters: (1) The maximum distance between two consecutive localisations that can be linked to the same particles, $R_{max}(\delta t, D)$, where $\delta t$ is the time interval between the frames and $D$ is the diffusion coefficient. (2) A memory parameter, that allows to fill gaps in individual trajectories owing to particles that transiently disappear below the detection limit or owing to fitting errors. (3) A minimum contrast value, that corresponds to the minimum particle mass that can be sent to the linking procedure. This parameter is mainly enabling us to avoid sending false positive localisations to the linking procedure due to the noise level. Here, $R_{max}\left(\delta t = \frac{1}{270Hz}, D_{max}\right)$ was set according to(*7*),

$$R_{max}(\delta t, D) \sim 2.55 \times \sqrt{4D\delta t}$$

where we considered a maximum diffusion coefficient of $1.5 \frac{\mu m^2}{s}$ that corresponds to the fastest particles on average and $\delta t$ was set to the inverse of the frame rate. This results in a maximum step size of 0.380 $\mu m^2$ that corresponds to 5 pixels. The memory parameter was set to one frame since our spike trimers have relatively high contrast. The minimum contrast value to send to the linking procedure was set here to -0.004 which corresponds to 110 kDa, close to our expected quantitative detection limit on bilayers.

Analysing trajectories to extract particle mass and diffusion coefficient

Detected trajectories where then screened similarly to the description in (*4*). Here we considered all trajectories that were longer than 111 ms (corresponding to 30 frames). The contrast histogram for each trajectory that was longer than 111ms was then examined by calculating the ratio between its median and average values to avoid trajectories where two different particles were combined in a single trajectory. Here, a threshold of 0.2 was set, which corresponds to a difference of more than 110 kDa between the median and the average of the trajectory as the smallest species we are examining are spike trimers with a mass of 550 kDa. Histograms were then fitted to a Gaussian function to extract the average contrast of the trajectory and its standard deviation. Poorly linked or fitted trajectories resulting in a high standard deviation that exceeded our calibrated standard deviation trend from landing and simulation were excluded. The fraction of these trajectories was around a few precent, and maxed out at 15% for very dense movies. The following procedure provides sufficiently long enough trajectories to obtain sufficient resolution and accuracy in mass and diffusion, while providing substantial statistics to correctly sample the distribution of different oligometric species as seen by comparing the analysis results with a simulated distribution.

Generating 2D Mass-Diffusion plots

Each detected trajectory that was analysed to extract the mass of a particle was also examined to calculate its diffusion coefficient. A spatial trajectory is defined by a set of positions $\{\boldsymbol{r}_{t_i}\}$, where $\boldsymbol{r}_{t_i}$ is the two-dimensional position of a particle at time point $t_i$. We calculated the set of distances, $\Delta r = \left\|\boldsymbol{r}_{t_{i+1}} - \boldsymbol{r}_{t_{i+1}}\right\|_2$ the particle has traveled between consecutive frames (or during the corresponding lag time, $t_{lag}$). A cumulative distribution was than calculated for the set of distances given by $P(\Delta r \leq r; t_{tag})$. For a particle diffusing on a bilayer undergoing Brownian motion, the diffusion coefficient was estimated as before (*4*) according to,

$$P\left[\Delta r \leq r; t_{\mathrm{lag}}\right] = 1 - exp\left(-\frac{r^2}{4D_{\mathrm{obs}}t_{\mathrm{lag}}+2\sigma^2}\right) \ ,$$

where, $\boldsymbol{\sigma}$ is the two-dimensional symmetric localisation error, and $\boldsymbol{D}_{\mathrm{obs}}$ is the effective observed diffusion coefficient that incorporates the fact that our measurement time resolution is slower than the real timescale of the Brownian motion of the particle on the supported lipid bilayer. Taking this into consideration, we assume the following expression for the observed diffusion coefficient,

$$D_{\mathrm{obs}} = D\left(1 - R\frac{t_0}{t_{\mathrm{lag}}}\right)$$

where $D$ is the Brownian diffusion coefficient, $t_0$ is the exposure time per frame, $R\frac{t_0}{t_{\mathrm{lag}}}$ is a correction for the diffusion coefficient arising from motion blur due to the finite measurement temporal resolution of a fast-moving Brownian particle, and the factor R depends on the acquisition parameters (*8*). In our experiments $t_0 \approx t_{\mathrm{lag}}$ and $R = 1/3$. The localization precision of our measurements as a function of mass and acquisition frame rate was estimated from simulations as described and shown in (*4*), where simulations where also used for validation of the value of $R$. Using these expressions, relating the experimental cumulative distribution of distances for individual tracked particles we can determine for each detected trajectory a corresponding diffusion coefficient, $D$. Given the extracted diffusion coefficient, the averaged trajectories contrasts can be corrected owing to motion blur that smears the detected psf and therefore lowers the fitted contrast by a few percent, depending on the mobility of the particle. Motion blur correction was validated both experimentally and with simulations for different masses, diffusion coefficients and acquisition parameters as described in (*4*) .

Correcting for contrast changes owing to optical path differences between the glass surface and spike on a supported lipid bilayer

When examining the dimensions of the soluble spike construct, we find that it can extend up to 30 nm away from the surface, when it is perfectly aligned perpendicular to the bilayer surface (**Fig S5a**). This extended structure can lead to a change in the measured interferometric contrast of spike when tethered through its his-tag to the bilayer. The change in the measured contrast is due to the added optical path difference between the reference light, reflected light from the water-glass interface and the scattered light from spike. To account for this variation for each data set, we measured this change by measuring spike trimer several times on glass and on the supported lipid bilayer and their contrast was then converted into mass using a protein standard for contrast to mass conversion as for all solution measurements. Measurements were done several times during each day of experiments, using the same mass photometer with the same acquisition parameters. The ratio between the two values for each spike contrast was recorded and used for correcting the mass of the detected species on the supported lipid bilayers (**Fig. S5d**). To validate that the origin of this difference in the measured interferometric contrast is indeed the extended shape of spike and not any bilayer related parameter, we performed a measurement where both spike and the standard protein were bound to the bilayer by their his-tag modification (**Fig. S5 b,c**). The standard protein forms oligomers in solution (**Fig S5c**) where its hexamer is measured at 540 kDa very close to the 533 kD measured for spike HexaPro (**Fig 5c**). When put together on the supported lipid bilayer (**Fig. 5b**), the standard protein mass does not change (since its center of mass is very close to the surface of the bilayer) as seen by the positions of the dimer, tetramer, and hexamer peaks (180, 360 and 540 kDa). However, the measured mass of spike was shifted to a lower mass owing to its different structure. This experiment with an internal control shows that the unique size and shape of the spike construct changes its measured contrast when tethered to the bilayer.

Thermodynamic model for 2D antibody induced spike oligomerisation on supported lipid bilayers

To model the distribution of oligomers on the supported lipid bilayer, we used the following thermodynamic framework. To develop the expression for the two-dimensional mole fractions, we assume the following: (1) Following incubation for about 1 h at room temperature the system is at steady state, and therefore the distribution of assemblies can be approximated by an equilibrium distribution. (2) The supported lipid bilayer also serves as a passivation layer, confining the ligands (ACE2/antibody) to the solution. In addition, the number of ligand molecules in solution ($N_l$) is much higher than the number of receptors (spike, $N_r$) on the bilayers ($N_l$ >> $N_r$), therefore, we can assume that the concentration of ligands in solution does not depend on binding to the receptor on the surface of the bilayer and is constant throughout the experiment. (3) Spike trimers are confined to the supported lipid bilayer interface and therefore the oligomerization reaction takes place on the membrane surface. (4) The concentrations of ligand in solution (≤ 150 nM) and the densities of spike on the bilayer (≤ 2 $\mu m^{-2}$) can be considered as a dilute solution or surface density.

Given these assumptions, the total Helmholtz free energy for a mixture of soluble ligand and tethered Spike proteins is given by the sum of the solution and surface free energies,

$$F_{\text{total}} = f_{\text{solution}} + f_{\text{surface}}. \qquad (S1)$$

Since spike trimers are confined to the bilayer by their three hexa-histidine tags, the first term, $f_{\text{solution}}$, is merely the free energy of soluble ligand and is given by,

$$f_{\text{solution}} = N_{\text{Av}}^{-1} V c_{\text{l}} ( \Delta f_{\text{l}}^{\circ} + k_{\text{B}} T \ln(c_{\text{l}} \cdot c_{\circ}^{-1}) - k_{\text{B}} T) \qquad (S2)$$

where, $N_{\text{Av}}$ and $V$ are Avogadro's number and volume, $c_l$ is the molar concentration of ligand molecules, $c_{\circ}$ is the reference concentration taking into account the concentration of ligands and water molecules (in the dilute solution taken as 55.5 M), $\Delta f_l^{\circ}$ is the standard change in free energy for the soluble ligand, $k_{\mathrm{B}}$ is Boltzmann constant and $T$ is the temperature. In a similar manner, we can write the free energy of receptors and ligands on the surface of the supported lipid bilayer:

$$f_{\text{surface}} = A \sum_{n_r,n_l} \frac{\rho_{n_r,n_l}}{n_r}\left(\Delta f_{n_r,n_l} + k_{\mathrm{B}}T \ln\left(\frac{\rho_{n_r,n_l}}{n_r} \cdot a_\circ\right) - k_{\mathrm{B}}T\right). \qquad (S3)$$

Here, A is a unit area, $\rho_{n_r,n_l}$ the number density of receptors in an assembled complex containing $n_r$ receptors and $n_l$ ligands, $\Delta f_{n_r,n_l}$ is the free energy change of assembling $n_r$ receptors and $n_l$ ligands into one complex, and $a_\circ$ is the cross-section area of one receptor. Since we assume constant concentration of ligand in solution, we can assume that the reaction on the surface proceeds at a constant chemical potential of free ligands. In this case, we can write the chemical potential of the ligand as,

$$\mu_{\mathrm{L}} = Const. = \left(\frac{\partial f_{solution}}{\partial c_l} \cdot \frac{N_{\mathrm{Av}}}{V}\right)_{V,T} = \mu_L^\circ + k_{\mathrm{B}}T \ln(c_l \cdot c_\circ^{-1}) \qquad (S4)$$

where $\mu_L^\circ$ is the standard chemical potential of the ligands and $c_l \cdot c_\circ^{-1} = X_l$ is their molar fraction. We can now use the result in **Eq. S4** to write the free energy term, $\Delta f_{n_r,n_l}$ in **Eq. S3**. This term corresponds to the free energy change of forming a complex of $n_r$ receptors and $n_l$ ligands. Since the driving force for forming the complex originates from the binding of ligand molecules, where $n_l$ must satisfy $n_l \geq n_r - 1$, the chemical potential of ligand must be considered in this term. Given this description and that all binding sites on the receptors are identical, independent and distinguishable, we can write $\Delta f_{n_r,n_l}$ under a fixed chemical potential of ligands in the solution as,

$$\Delta f_{n_r,n_l} = -k_{\mathrm{B}}T \ln\left(q_{n_r,n_l}\lambda_{\mathrm{L}}^{n_l}\right) = -k_{\mathrm{B}}T \ln\left(\frac{\Omega_{n_r,n_l}}{n_r!\,n_l!} e^{-\frac{\epsilon(n_l)}{k_{\mathrm{B}}T}} \lambda_{\mathrm{L}}^{n_l}\right). \qquad (S5)$$

In **Eq. S5**, $q_{n_r,n_l} = \frac{\Omega_{n_r,n_l}}{n_r!n_l!} e^{-\frac{\epsilon(n_l)}{k_{\mathrm{B}}T}}$ is the partition function for an $n_{r,l}$ complex. The first term $\frac{\Omega_{n_r,n_l}}{n_r!n_l!}$ corresponds to the configurational degeneracy in assembling the complex, with $\Omega_{n_r,n_l}$ the number of states to arrange $n_r$ receptors with $n_l$ ligands, and the two factorials were included to account for the fact that both the receptors molecules and the ligands are indistinguishable. The second term $e^{-\frac{\epsilon(n_l)}{k_{\mathrm{B}}T}}$, includes the free energy gain, $\epsilon(n_l)$, in binding $n_l$ ligands to form the complex. The last term, $\lambda_{\mathrm{L}}^{n_l}$, introduces the chemical potential of ligands in the solution, $\lambda_{\mathrm{L}}^{n_l} = e^{n_L \frac{\mu_L}{k_{\mathrm{B}}T}}$. Rearranging **Eq. S5** gives,

$$\Delta f_{n_r,n_l} = -k_{\mathrm{B}}T \ln\left(\frac{\omega_{n_r}}{n_r!}\right) - k_{\mathrm{B}}T \ln\left(\frac{\omega_{n_l;n_r}}{n_l!}\right) + \epsilon(n_l) - n_l\mu_L = n_l\Delta\tilde{\mu} - k_{\mathrm{B}}T \ln\left(\frac{\omega_{n_r}}{n_r!}\right), \qquad (S6)$$

where, $\omega_{n_r}$ and $\omega_{n_l;n_r}$ correspond to the number of possible arrangements of $n_r$ receptors and $n_l$ ligands on a complex that contains $n_r$ receptors, respectively. The last definition $\Delta\tilde{\mu} = \frac{\epsilon(n_l)}{n_l} - \frac{k_{\mathrm{B}}T}{n_l} \ln\left(\frac{\omega_{n_l;n_r}}{n_l!}\right) - \mu_L$, is the average change in chemical potential between the soluble ligand ($\mu_{\mathrm{L}}$) and a ligand that is bound to a complex containing $n_r$ receptors and $n_l$ ligands. This term depends on the details of the interaction between the ligand and the receptor and will be derived for each binding pair. The second term on the right-hand side of **Eq. S6** is identical for all binding partners and corresponds to the configurational entropy of $n_r$ multivalent subunits (in this case the receptors) that assemble into one complex. Here, the configurational entropy of the receptors was calculated as previously suggested by Goldberg (*9*) for the case of branched oligomers of multivalent subunits without considering the possibility of ring formation. Under these conditions the number of microscopic states, $\omega_{n_r}$, for receptors of valency $\nu$, is given by,

$$\omega_{n_r;\nu} = \frac{(\nu n_r - n_r)!\,\nu^{n_r}}{(\nu n_r - 2n_r + 2)!}. \qquad (S7)$$

Finally, taking into account the fact that the receptors are indistinguishable, we divided $\omega_{n_r}$ by $n_r!$ to give,

$$\Omega_{n_r} = \frac{\omega_{n_r}}{n_r!} = \frac{(\nu n_r - n_r)!\,\nu^{n_r}}{(\nu n_r - 2n_r + 2)!\,n_r!}. \qquad (S8)$$

To derive the equilibrium oligomeric distribution of the receptors-ligand complexes on the supported lipid bilayer, $\rho_{n_r,n_l}$, we minimised $f_{\text{surface}}$ (**Eq. S3**) subject to the constraint of receptor mass conservation,

$$\sum_{n_r} \rho_{n_r} = \rho_{total}. \qquad (S9)$$

The equilibrium distribution is therefore given by solving the following equation using the Lagrange multipliers method:

$$\frac{\partial}{\partial \rho_{r,l}}\left(f_{\text{surface}} - \lambda\left(\sum_{n_r} \rho_{n_r} - \rho_{total}\right)\right) = 0, \qquad (S10)$$

which results with the distribution,

$$\rho_{r,l} = \rho_\circ n_r \exp\left(-\frac{\Delta f_{n_R,n_l} - n_r\mu_R}{k_B T}\right) = \rho_\circ n_r \exp\left(-\frac{n_l\Delta\tilde{\mu} - k_B T\ln\left(\frac{\omega_{n_r}}{n_r!}\right) - n_r\mu_R}{k_B T}\right), \qquad (S11)$$

where $\rho_\circ$, the reference state was set to $\rho_\circ = a_\circ^{-1}$, and the Lagrange multiple, $\lambda$, was replaced by the corresponding chemical potential of free receptor,

$$\mu_R = \left(\frac{\partial f_{\text{surface}}}{\partial \rho_{n_r,n_l}} A^{-1}\right)_{V,T} = \mu_R^\circ + k_B T\ln(\rho\cdot\rho_\circ^{-1}). \qquad (S12)$$

<u>Interaction model for the spike-antibody system</u>

To obtain the distribution of surface oligomers for different spike-ligand interaction systems and concentrations (**Fig. 4 and S13**), we used the following expression to account for the interaction term $\Delta\tilde{\mu}$,

$$\Delta\tilde{\mu} = n_L^{-1}\big((n_L - n_r + 1)\epsilon_1 + (n_r - 1)\epsilon_2\big) + k_B T\ln\big(\Omega_{nL,n_r}\big) - \mu_L^\circ - k_B T\ln(c_L\cdot c_\circ^{-1}). \quad (S13)$$

In **Eq. S13** the first parameter, $\epsilon_1$, is the effective free energy change when a ligand binds to one binding site of spike trimer, and $\epsilon_2$ is the free energy change in the binding of a ligand that results in oligomerization of two spike trimers (ligand that binds two binding sites from different spike trimers). The third term is the configurational degeneracy of the ligands that are bound in a complex of size $n_r$ spikes, where $\Omega_{n_l,n_r} = \frac{\omega_{n_l,n_r}}{n_l!}$. The last two terms correspond to the chemical potential of the ligands in solution. **Eq. S13** can be rearranged as,

$$\Delta\tilde{\mu} = \Delta\tilde{\mu}^{\circ} + n_L^{-1} k_{\mathrm{B}} T \ln\left(\Omega_{nL,n_r}\right) - k_{\mathrm{B}} T \ln\left(c_L \cdot c_{\circ}^{-1}\right), \qquad (S14)$$

where $\Delta\tilde{\mu}^{\circ}$ is the change in the standard chemical potential between soluble ligand and ligand that is bound in a complex, and the last two terms correspond to the entropy change between these two states.

<u>Parameters for interaction of antibodies 159, 384 and 150 with spike</u>

*Antibody 159* binds the NTD domain of spike (*10*) and therefore can occupy up to three binding sites per trimer (**Fig. 4 a,b and S11**). Since the NTD binding sites are separated by a distance larger than the distance between adjacent RBD domains, and there is no known conformational state that facilitates preferred binding (such as "down" and "up" states for the RBD) we expect little or no cooperativity between the different NTD binding sites for COVOX159. Here, the model included the two free energy changes, $\epsilon_1$, for antibody binding to one site, and the oligomerisation interaction, $\epsilon_2$, calculated by,

$$\epsilon_2 = \epsilon_2^{\circ} + \Delta(n_r - 2), \qquad (S15)$$

where the parameter $\Delta$ was added to include a possible small crosslinking penalty that may arise by the need to assemble increasingly larger, closely packed spike trimers at specific orientation compared with their anchoring point to the lipid bilayer. The fitted values of the three parameters are summarised in **Table S2**. The configurational degeneracy $\Omega_{n_l,n_r}$ was calculated by the number of possible arrangements to bind $n_l$ indistinguishable bivalent ligands to $\nu n_r$ multivalent receptors (for spike $\nu = 3$) binding sites, while $n_r - 1$ of the ligand and $n_r$ of the binding sites are involved in oligomerisation interactions (similar to the calculation presented in (*9*)). This results in the following expression for the configurational degeneracy:

$$\Omega_{n_l,n_r} = \binom{(\nu-2)n_r + 2}{n_l - n_r + 1} \cdot 2^{n_l} = \frac{\left((\nu-2)n_r + 2\right)!}{(n_l - n_r + 1)!\left((\nu-1)n_r + 1 - n_l\right)!} \cdot 2^{n_l}. \qquad (S16)$$

*Antibody 384* binds the RBD of spike, preferably to its "down" conformation as reported previously (*10*) and also shown here by its improved binding to the HaxaPro stabilised spike (**Fig. 4f and S12**), which maintains the RBD in its "down" conformation (*11*). Since the distance between adjacent RBDs is relatively short, we expect cooperativity between the binding sites, similarly to the observed binding of ACE2. Indeed, although the antibody binds spike with a $K_{\mathrm{D}} \approx 1$ nM no more than two antibodies were observed simultaneously on the same spike trimer. Similarly, we observed no more than 4 antibodies binding to a crosslinked aggregate of two spike trimers. To account for cooperativity in a similar way to the interaction in solution involving

ACE2, we included a cooperativity parameter to the binding of antibodies from solution to tethered spike. The free energy change for binding from solution, $\epsilon_1$ was given as,

$$\epsilon_1 = \epsilon_1^{\circ} + \alpha n_l', \qquad (S17)$$

where $n_l'$ is the number of prebound ligands on a spike trimer prior to the specified binding event. For example, for the freely diffusing spike trimer, for the first binding event, the free energy gain was set to $\epsilon_1^{\circ}$, the second binding event results in a free energy gain of $\epsilon_1^{\circ} + \alpha$, and the third binding site with $\epsilon_1^{\circ} + 2\alpha$. A preformed crosslinked complex (one antibody already bound to 2 RBDs from two different spike trimers) was considered as a "new spike" receptor with 4 available binding sites. In this case the first binding event from solution will result in a free energy of $\epsilon_1^{\circ}$, while the second one to the same spike trimer will result in $\epsilon_1^{\circ} + 2\alpha$ as this is the third RBD to be occupied on the same trimer. For the configurational degeneracy of the crosslinked structures, $\Omega_{n_l,n_r}$, we accounted for the fact that there are two different positions with different energetics (see **Fig S18**). For the first two binding events (binding to sites 1) the degeneracy factor was calculated as $\Omega_{n_l,n_r} = 4 \cdot 2^{n_l}$. For the remaining binding events, which must bind sites of type 2 we calculate the degeneracy as,

$$\Omega_{n_l,n_r} = \binom{(\nu-2)n_r + 2}{n_l - n_r + 1} \cdot 2^{n_l} = \frac{\big((\nu-2)n_r + 2\big)!}{(n_l - n_r + 1)!\,\big((\nu-1)n_r + 1 - n_l\big)!} \cdot 2^{n_l}. \qquad (S18)$$

*Antibody 150* has a unique binding pattern. While its affinity to spike trimer ($K_D$ ≈1 nM) is similar to 384, its oligomerisation propensity is very low. In addition, the binding of the first antibody seems to induce very strong negative cooperativity for further binding. This could be explained by the fact that the first binder partially blocks the adjacent RBD, or that it affects the probability of the adjacent RBD to be found in the "up" conformation which enables the binding of 150. We introduce a negative cooperativity term as for antibody 384 (**Eq. S17**). Since the induced oligomerisation is very weak, we could not determine whether the free energy of oligomerisation ($\epsilon_2$) decreases with the size of the oligomer, therefore it was neglected here and we set $\epsilon_2 = \epsilon_2^{\circ}$. The configurational degeneracy was taken to be similar to antibody 384, however owing to the strong negative cooperativity and low oligomerisation propensity, which results in a small amount of oligomers and its significance in term of relative mole fraction is low.

Comparison between monovalent and divalent binding curves in Figure 5

To generate the expected binding curves at different surface densities (**Fig. 5**) we calculated the expected thermodynamic distribution for three cases: (1) Monovalent antibody binding to spike on the surface (taking the interaction parameters of COVOX159 as the expected affinity of the Fab domain), (2) bivalent antibody COVOX159 in solution and spike on the surface, (3) ACE2 in solution and spike on the surface, and (4) spike in solution and ACE2 on the surface.

(1) For the monovalent case the following model was assumed, where we defined $P_n$ as the probability of a spike trimer to be bound to $n$ ligands, with $P_n$ defined by,

$$P_n = \sum_{n=0}^{M} \frac{\rho_n}{\rho_{\text{total}}} \ , \qquad (S19)$$

where $\rho_{\text{total}}$ is the initial surface density of the receptors on the surface and $\rho_n$ is the surface density of receptors bound to $n$ ligands. In the case of a monovalent ligand, the surface density of receptors does not play a role and the occupancy of the ligand on the receptors can be described on the basis of the Langmuir isotherm. In this case, the free energy of interaction between $n$ ligands and a receptor is given by,

$$\Delta G = n \cdot \Delta\mu + k_{\text{B}}T \ln \Omega_{M,n}\,, \qquad (S20)$$

where $\Delta\mu$ is the change in chemical potential upon binding of a single lignad to a single binding site on the receptor assuming a constant concentration of ligand in the solution and is given similarly to **Eq. S4**. The second term in **Eq. S20** again represents the configurational entropy, where $\Omega_{M,n} = \binom{M}{n} = \frac{M!}{n!(M-n)!}$. The average occupancy fraction, $\Theta$, for monovalent ligands is therefore,

$$\theta(C_{\text{L}}) = \frac{\overline{n(C_{\text{L}})}}{M} = \frac{1}{M}\sum_{n=0}^{M} n \cdot P_n = \frac{1}{M \cdot Z}\sum_{n=0}^{M} n \cdot exp(-\Delta G_n/k_{\text{B}}T), \qquad (S21)$$

where $Z$ is the partition function and is equal to $\sum_{n=0}^{M} \exp -\Delta G_n/k_{\text{B}}T$. The interaction strength of a single ligand to a single binding site on spike trimer, $\Delta\mu$ was set to 21.29 $k_{\text{B}}$T similar to the interaction parameter of antibody COVOX159.
(2) For the bivalent case of antibody binding to spike, we used the same model as previously described for the case of COVOX159 binding while changing the total density of spike on the surface.
(3) For the bivalent case of ACE2 binding to spike, we used the same model as previously described for the case of COVOX384, however, we changed the interaction free energy to match the experimental values as described in Table S1 for the interaction between wtSpike and ACE2 from solution.
(4) For the trivalent case of spike binding to a membrane surface containing ACE2, we used the parameters of the interaction between omSpike and ACE2 that includes a maximum of two occupied binding sites per spike with positive cooperativity for the second binding site. This allows formation of linear oligomers.

Extracting relative abundances and fitting thermodynamic distributions

Mole fractions of the different complexes formed by the spike-antibodies interactions were determine by fitting a set of Gaussian functions to the measured histograms (**Figs. S10-12**). The Gaussian functions were constrained to the expected mass of the different complexes and allowed to deviate from these values by 2-4%. The widths of the peaks were also constrained to the expected mass resolution <50 kDa. Following peak assignment and fitting, the mole fractions of spike in oligomers of size $n_{spike}$ were calculated by multiplying the relative areas by the number of spiks trimers in the complex and then the distribution was normalised. Following the determination of the mole fraction for each spike-antibody pair (containing 3 antibody concentrations) a global fit yielded the 2D and 3D interactions standard free energy changes (**Fig. 13**). The global fit was performed by a custom-written python script using the Scipy optimisation package. The extracted free energy changes were used to calculate the 2D and 3D dissociation constants.

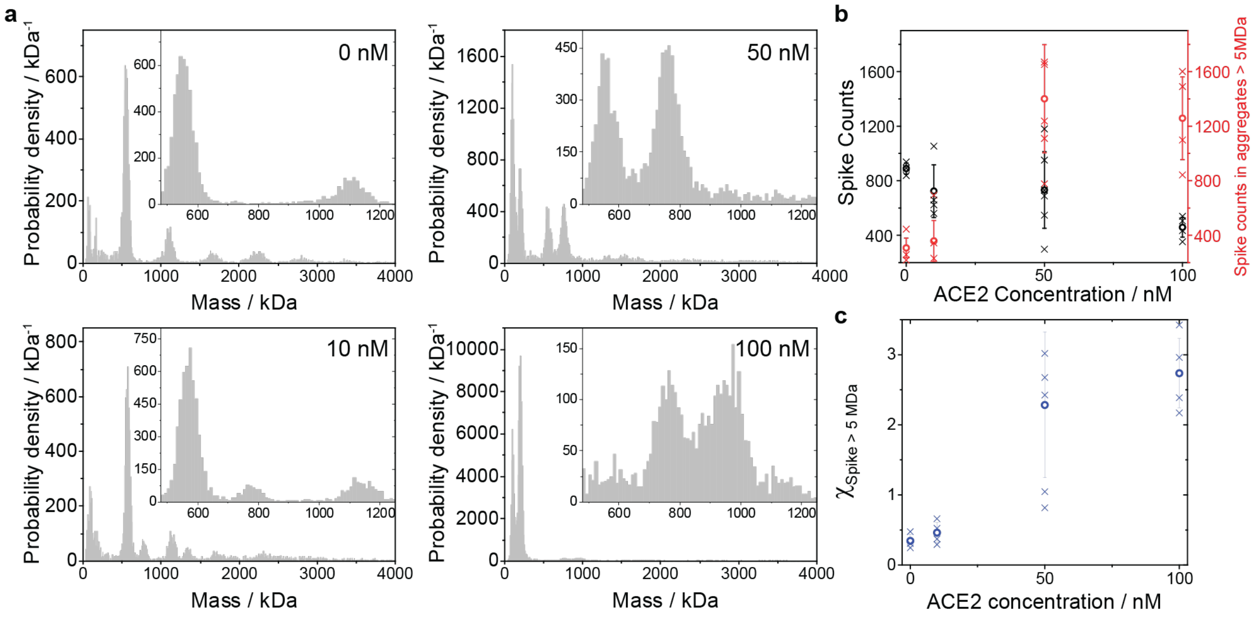


**Fig. S1 ACE2-Spike interaction at µM concentration in solution.**

(a) Mass histograms at a final concentration of 50 nM spike with 0 to 100 nM ACE2 following our rapid dilution protocol from µM concentration (see Materials and Methods). Each histogram contains the cumulative counts from 3 - 6 technical replicates. All histograms were corrected to remove small mass impurities as explained and shown in **Fig. S16**. (b) Number of counts for molecular species containing only one spike trimer (black symbols and left y-axis) and estimated number of spike trimers in molecular species of mass larger than 5 MDa (red symbols and right y-axis). The estimated counts were based on the mass of the detected complex divided by the mass of the spike trimer bound to one ACE2 dimer, which form the minimal subunit for oligomerisation. Crosses correspond to individual technical repeats, while circles and error bars correspond to the averages and standard deviations. (c) Relative number of counts in oligomers larger than 5 MDa compared to the number of counts of complexes containing only one spike trimer as a function of ACE2 concentration.

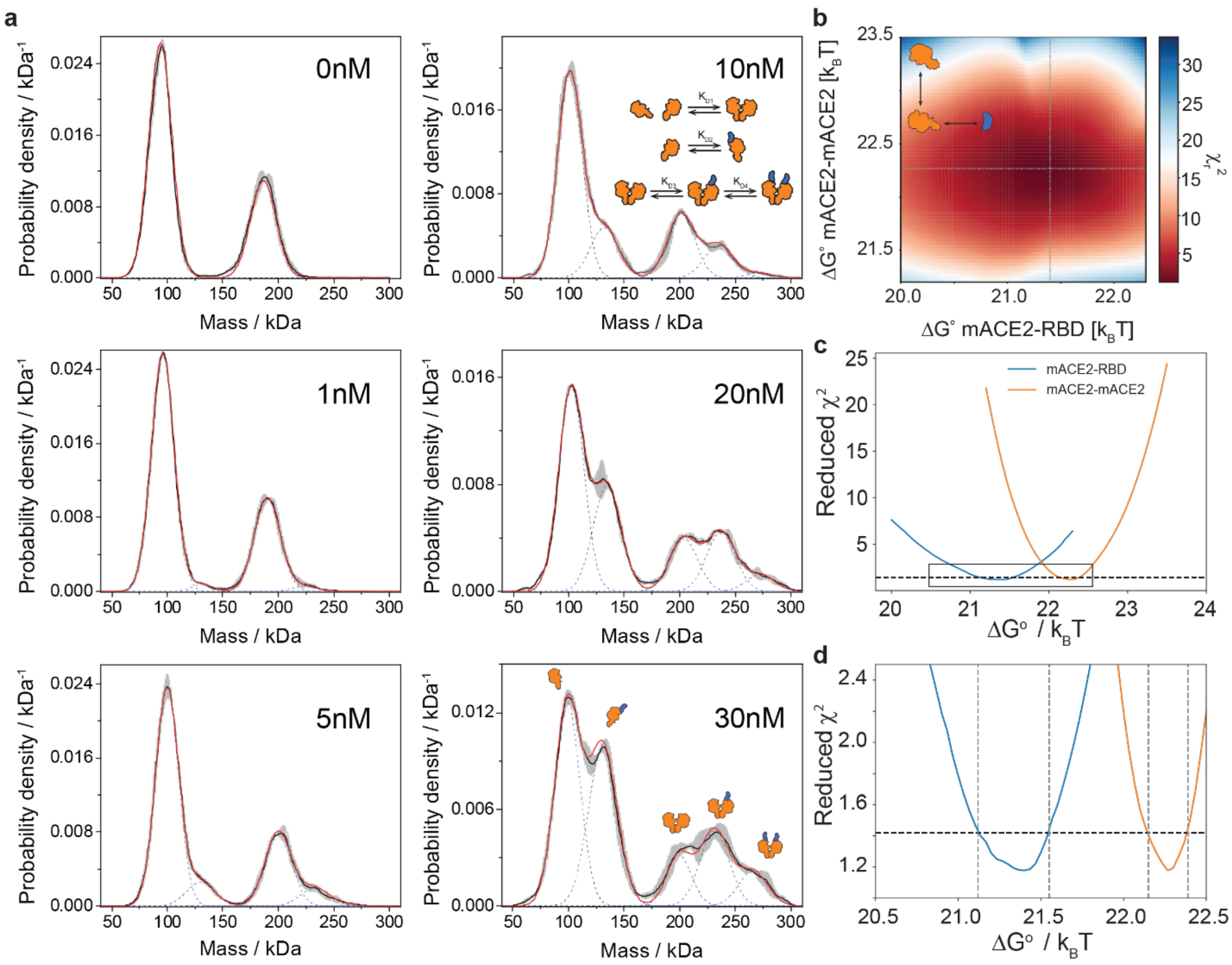


**Fig. S2. Mass distributions and global fit analysis of the ACE2–RBD interaction.**

(a) Fitting results for measured 10 nM ACE2 with a range of RBD concentrations (0-30 nM). Black curves and grey error bars correspond to the average probability density plot and standard deviations of three technical replicates, blue broken lines correspond to the individual Gaussian peaks used to generate the best fit, indicated by the red curve (see Material and Methods). Panels (b) and (c), Confidence limit estimation for the values of the standard free energy changes ($\Delta G°$) associated with the interaction of two monomeric forms of ACE2 (orange curve) and with the interaction between the RBD and one monomeric ACE2 (bule curve). The value of the reduced $\chi^2$ function was plotted along the two lines indicated in Figure 2b, corresponding to the two free energy values that minimise the reduced $\chi^2$. Panel (d) shows a zoom of the range of values close to the global minima for each free energy parameter. Fitting errors were estimated from the expected standard deviation of the reduced $\chi^2$ function based on fitting six solution conditions simultaneously. The resultant value of $\frac{\sigma_{\chi^2}}{\sqrt{6}}$ was added to the minimal value of the reduced $\chi^2$ and the confidence limits were taken accordingly as plotted by the dashed lines in (d).

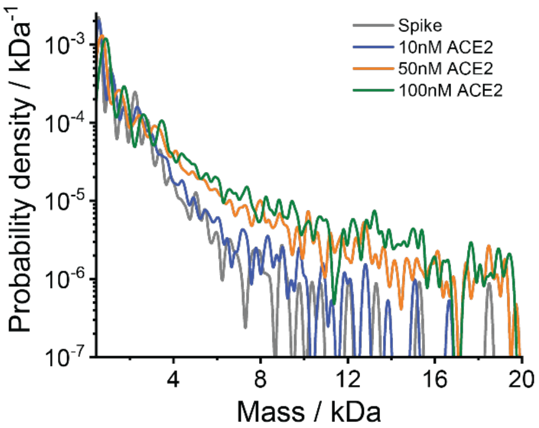


**Fig. S3. Oligomerisation of Spike as a function of ACE2 concentration.**

Average mass histograms plotted on a semilogarithmic scale for different mixing ratios of soluble wtSpike and ACE2. Spike trimer was mixed at concentrations of 0.55 μM with ACE2 concentrations ranging from 0.33 to 3.3 μM. Probability densities were computed using kernel density estimators with a bandwidth of 100 kDa. Only masses larger than 450 kDa were considered for probability calculation. Each probability density represents an average of 3 - 6 technical repeats.

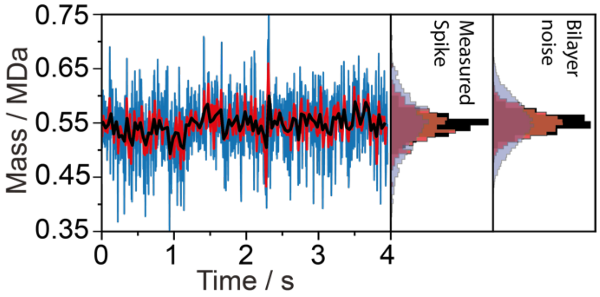


**Fig. S4. Mass fluctuations during single particle trajectories**

Continuous mass trace of a single spike trimer as it diffuses on a supported lipid bilayer (also shown in **Fig. 2c**). Blue curve corresponds to the trace as it was tracked and fitted at 270 Hz as used throughout our experiments. The red and black curves correspond to the same trace following the averaging of consecutive windows of 4 and 10 measurements along the trace, representing an effective frame rate of 67.5 and 27 Hz, respectively (without the effect of motion blurring). Middle panel shows the resultant histograms for the three traces. The right panel, shows the histogram of values from a single pixel at the center of the FOV (in mass units) from a background subtracted bilayer movie without adding spike. This histogram represents the intrinsic background fluctuations/noise level. The center of the histogram was shifted from zero to the average mass of the spike particle for clarity.

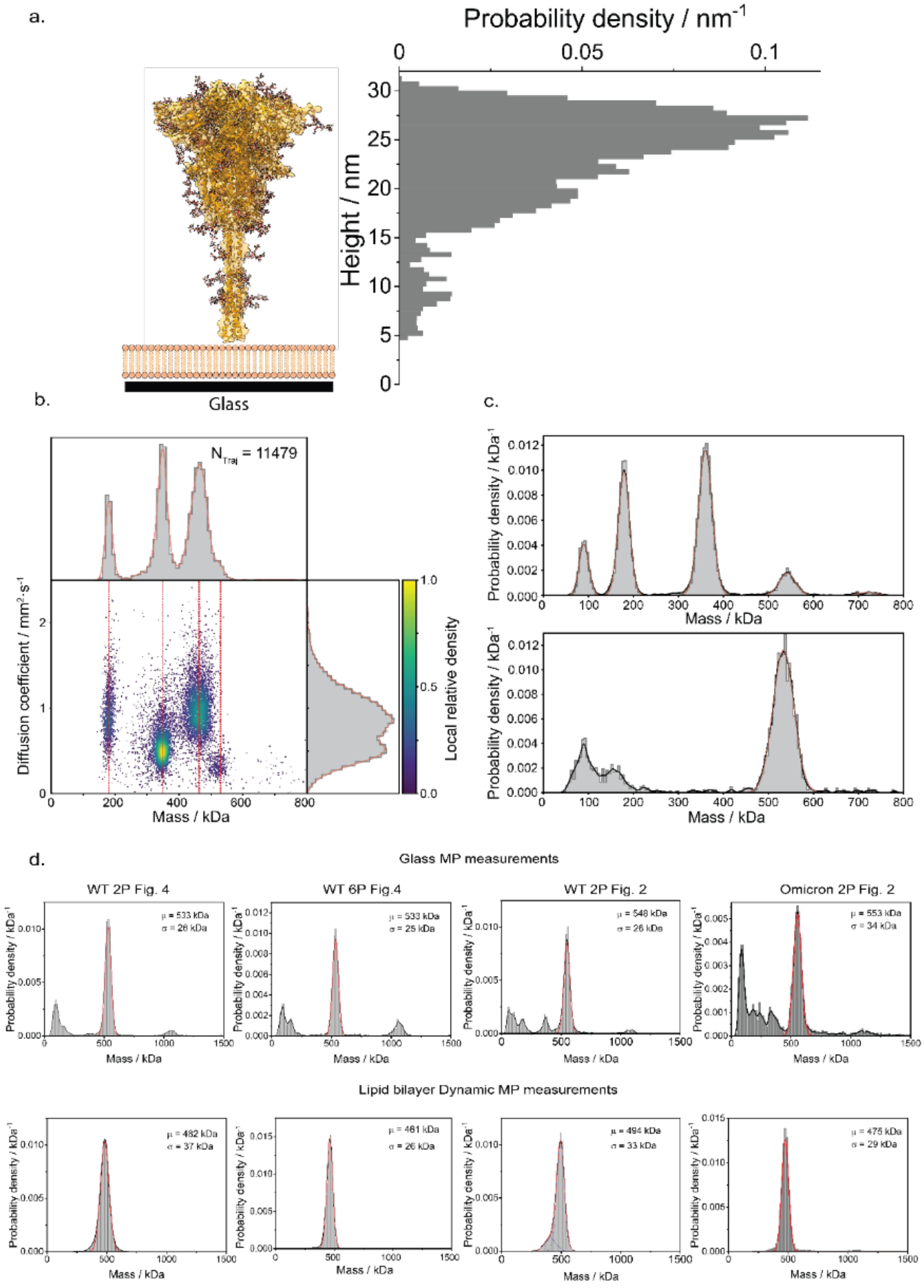


**Fig. S5. The effect of spike size and morphology on measured contrast**

(a) Atomic structure of spike trimer (based on PDB entry (*12*)) following insertion of missing residues and glycans by MD simulations (*13*). We removed the trans membrane domain to represent the structure of the soluble domain used in our study. The trimer was aligned to the z axis using PCA analysis and its mass density along z was plotted as a histogram with a bin size of 0.5 nm. An additional shift of 5 nm was included to represent the tethering on the supported lipid bilayer. (b) 2D dynamic MP plot of diffusion *vs* mass for simultaneously diffusing HexaPro stabilized spike with Dynamin 1 ΔPRD that was used as a protein mass standard. Both proteins are tethered to the bilayer by the interaction between the protein his-tag and lipid NTA(Ni). Red vertical lines correspond to the mass of the fitted species including for dimer, tetramer and hexamer of Dynamin (masses: 180, 350, and 530 kDa) and spike trimer (463 kDa). (c) Mass histogram of Dynamin (top) and spike (bottom) measured on a standard MP landing assay on a glass surface. Fitted peak masses are 90, 180, 360, 542, 720 kDa for dynamin monomer to octamer and 534 kDa for spike. (d) Comparison between the measured mass of spike using standard MP in solution and mass measured on supported lipid bilayers. The measurements show the different in mass for every data set presented in this work. For each data set we used the ratio between the two methods to correct bilayer-derived MP histograms for each data set.

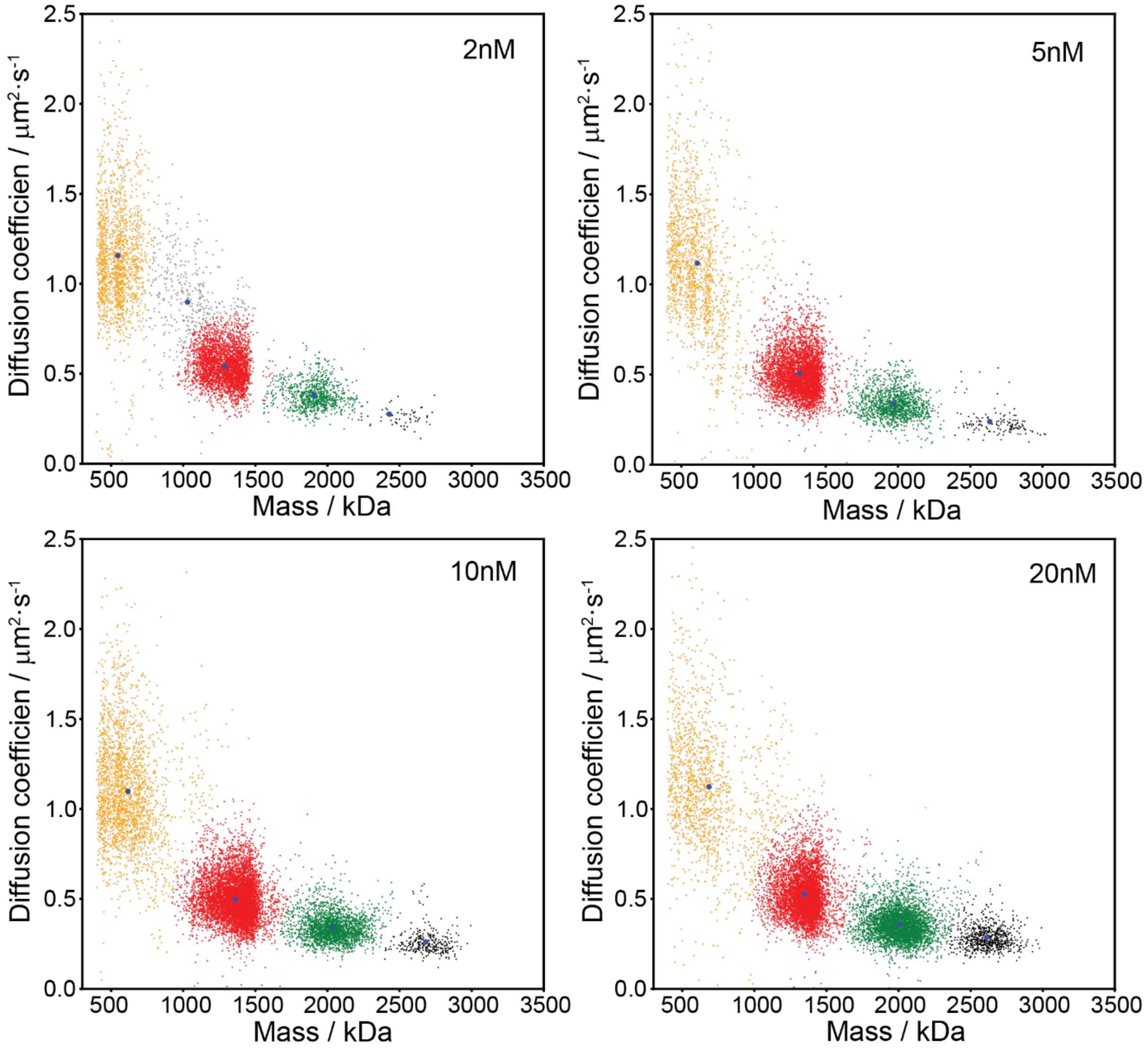


**Fig. S6. Dependence of diffusion coefficient on oligomer size.**

Representative data from each antibody-Spike concentration. Clustering was performed using the function *sklearn.mixture.BayesianGaussianMixture* from the scikit-learn python package. The number of clusters was either four or five to account for a possible need to achieve convergence depending on noise. For each cluster (colored orange, red green and black) the average and standard deviation were taken. The data in Fig. 2h show the results from four different antibody concentrations and two technical repeats, each.

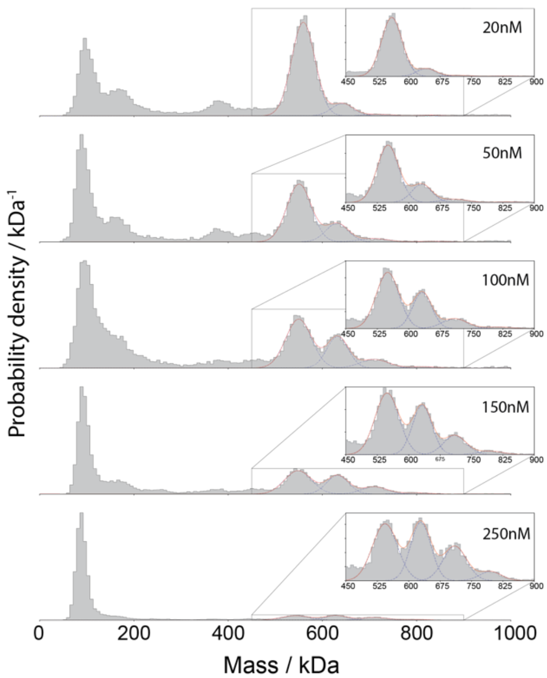


**Fig. S7. Quantification of the interaction between wtSpike and monomeric ACE2**

Normalised cumulative mass histograms following incubation of 25 nM wtSpike trimers with 20, 50, 100, 150 and 250 nM monomeric ACE2. Insets show the mass range between 450 and 900 kDa, where the resulting complexes containing wtSpike with 0, 1, 2 and 3 ACE2 are detected. The red curve corresponds to the sum of 4 Gaussian functions that were used to fit the histograms (see Materials and Methods). Dashed blue curves correspond to the individual fitted Gaussian functions. Bin size of the histograms are 5 kDa. Histograms correspond to the cumulative counts of all technical replicas, the relative probabilities shown in **Fig. 3d-f** were extracted by fitting the individual repeats.

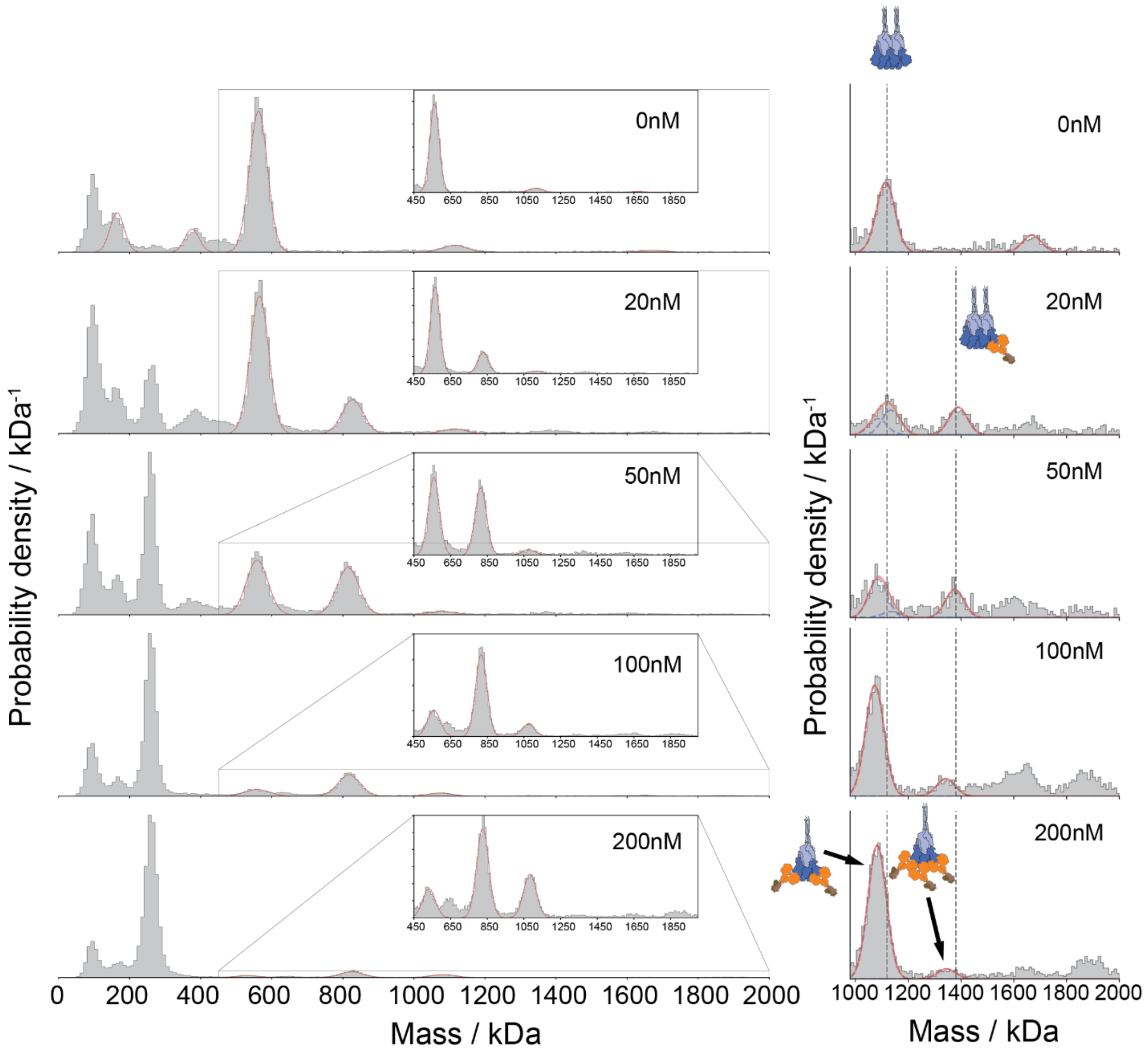


**Fig. S8. Quantification of the solution interaction of wtSpike with dimeric ACE2.**

Normalised cumulative mass histograms after mixing 25 nM wtSpike with 0, 20, 50, 100 and 200 nM dimeric ACE2. Insets show the mass range between 450 and 2000 kDa where the masses of wtSpike trimers, and dimers of trimers interacting with ACE2 are expected. Histograms were generated as described in **Fig. S5**. The top histogram shows the mass distribution of wtSpike without added ACE2, indicating a small number of dimers of trimers, as well as small amounts of spike dimers and monomers at the low mass range. The zoom on the right shows that increasing the ACE2 concentrations the abundance of spike with two ACE2 dimers bound becomes separated from the peak of spike dimers. Our results also suggest the existence of small amounts of spike bound to three ACE2, however the total number of counts is too small for a confident assignment.

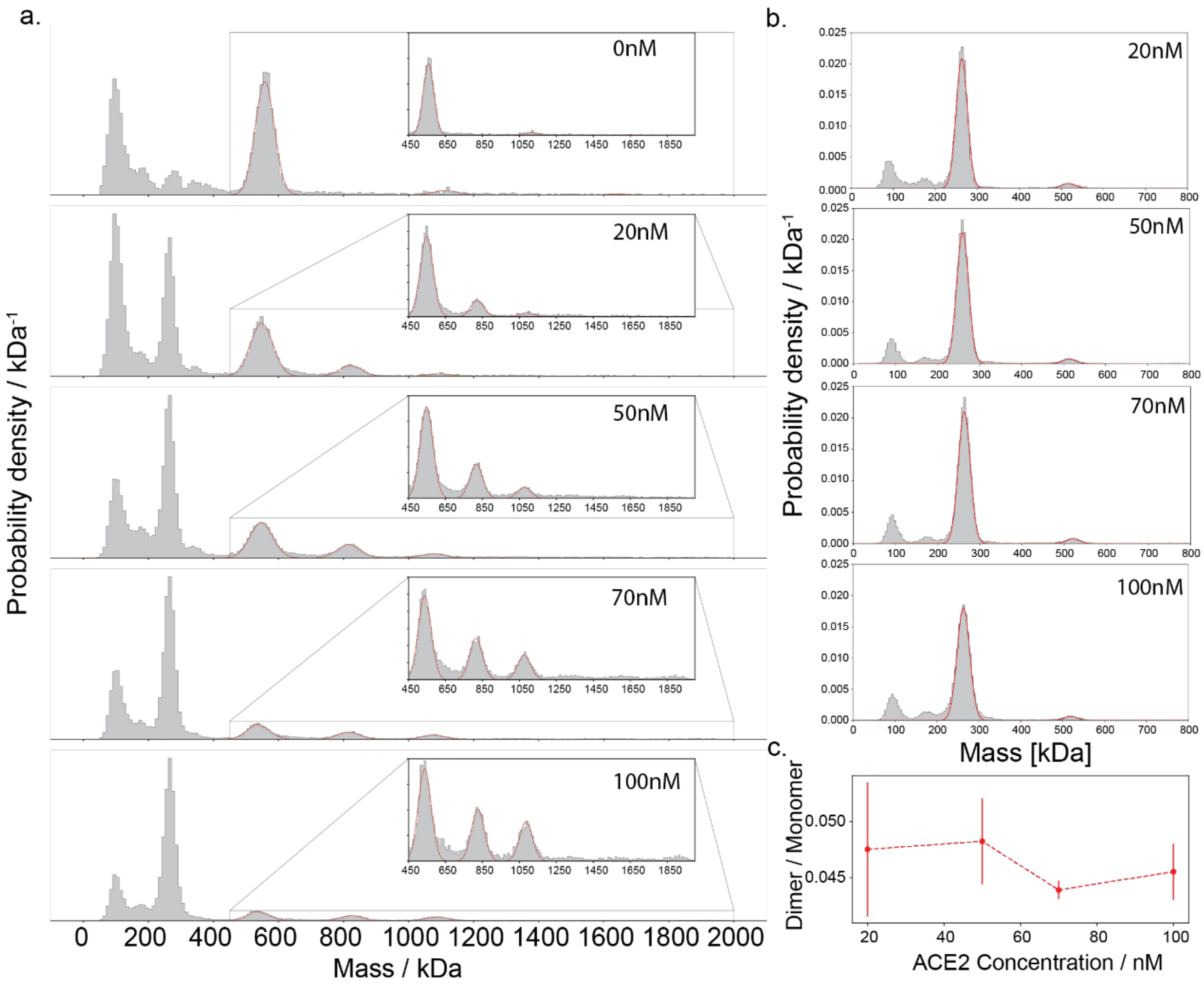


**Fig. S9. Quantification of the solution interaction of omSpike with dimeric ACE2.**

(a) Normalised cumulative mass histograms measured after mixing 25 nM omSpike with 0, 20, 50, 70 and 100 nM dimeric ACE2. Insets show the mass range between 450 and 2000 kDa where the masses of omSpike trimers, and dimers of trimers interacting with ACE2 are expected. Histograms were generated as described in **Fig. S5**. Top histogram shows the mass distribution of omSpike alone. (b) Normalised cumulative mass distribution of ACE2 dimers showing the small amount of possible ACE2 aggregates. The abundance ratio (c) was calculated by fitting Gaussians to the two peaks and dividing their fitted areas. The mass of ACE2 aggregates is close to the mass of spike trimer and the constant ratio of aggregates at different concentrations of ACE2 allows us to subtract the excess resulting counts from overlapping peaks of free spike by multiplying the counts of free ACE2 by the same factor. Owing to the small number of counts the excess counts will only have a small affect (10-20% if correction would not apply) on the relative abundance of free spike at high concentration (70 and 100 nM).

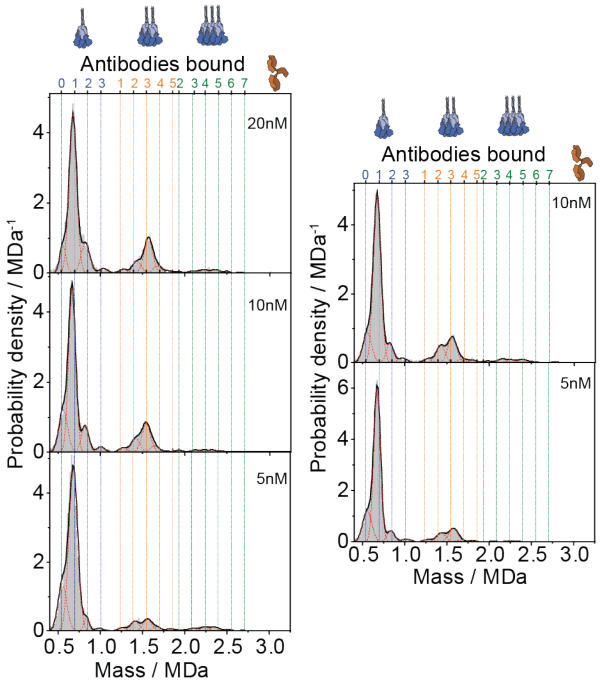


**Fig. S10. WtSpike interaction with COVOX150 antibody**

The histograms (grey) are the averaged measured masses of individual detected trajectories of wtSpike-COVOX150 complexes as they diffuse on the supported lipid bilayer. Each histogram was fitted to a sum of Gaussian functions constrained to the expected mass of the spike-antibody complex (see Materials and Methods), with a histogram bin size of 10 kDa. The black line corresponds to the sum of the Gaussians functions and the red dashed lines correspond to the individual fitted functions. Colored vertical lines show the expected masses of spike-antibody complexes with blue, orange and green corresponding to spike trimer, dimer of trimers and trimer of trimers, respectively. The number of antibodies in each complex is indicated by the numbers on the top of each panel with the respective colors. The two panels show independent repeats.

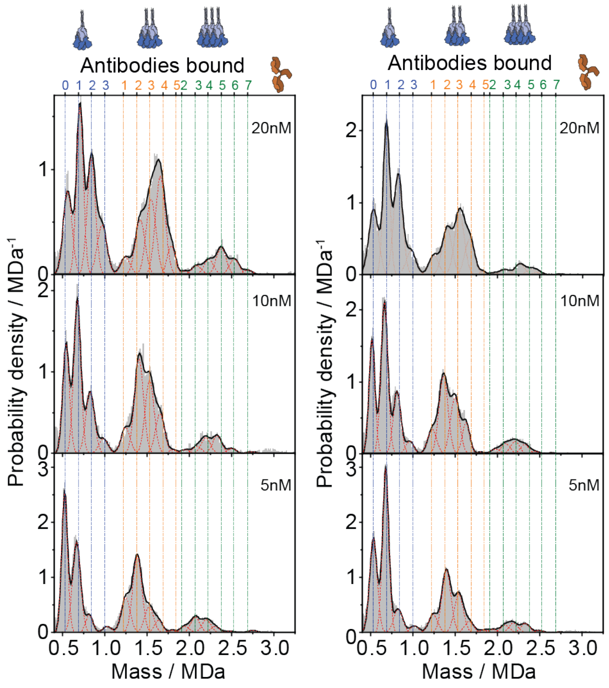


**Fig. S11. Binding of COVOX159 antibody to wtSpike**

Histograms and fits were generated as described for Fig. **S15** and materials and Methods. Colored vertical lines show the expected masses of spike-antibody complexes with blue, orange and green corresponding to spike trimer, dimer of trimers and trimer of trimers, respectively. The number of antibodies in each complex is indicated by the numbers on the top of each panel with the respective colors. The two panels represent two independent repeats for each antibody.

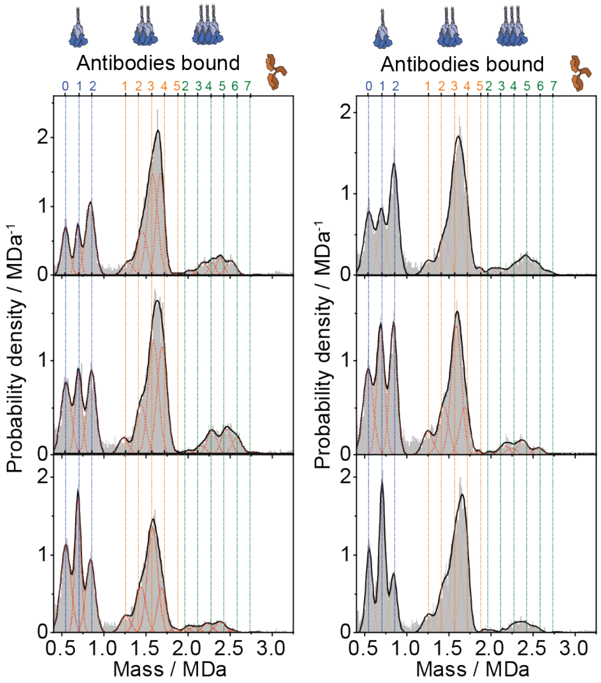


**Fig. S12. COVOX384 antibody binding to wtSpike**

Histograms and fits were generated as described in **Fig. S15** and Materials and Methods. Colored vertical lines show the expected masses of spike-antibody complexes with blue, orange and green corresponding to spike trimer, dimer of trimers and trimer of trimers, respectively. The number of antibodies in each complex is indicated by the numbers on the top of each panel with the respective colors. The COVOX384 antibody shows a maximum of 2 antibodies binding to a single spike. The two panels represent two independent repeats for each antibody concentration.

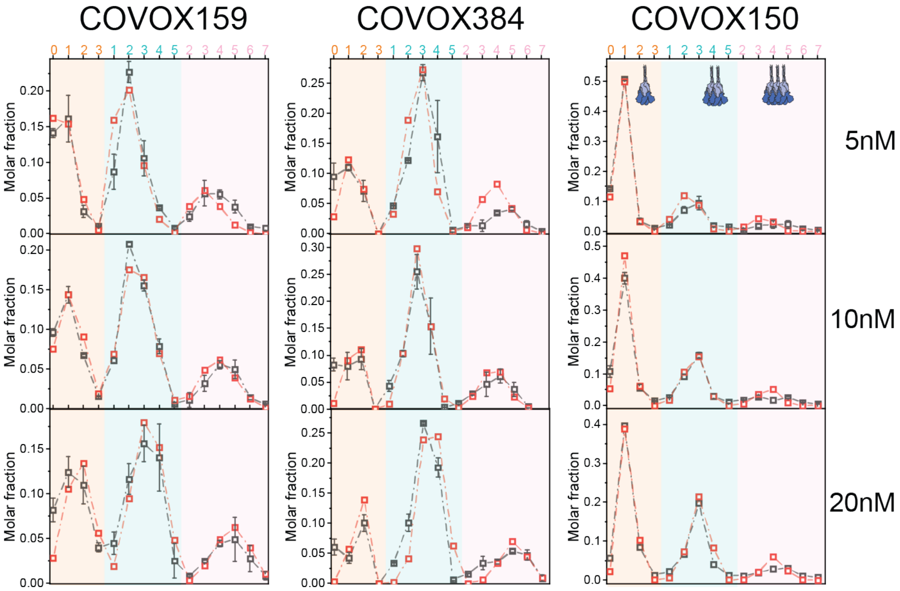


**Fig. S13. Determining 2D and 3D affinities from extracted mass fractions.**

Fitting results of the thermodynamic model to the surface spike mole fractions extracted from Dynamic-MP data. Grey symbols and error bars correspond to the experimentally extracted mole fractions of the different detected complexes on the surface of the bilayer (**Figs. S15-17** and Materials and Methods). Mole fractions were calculated by the relative areas of the fitted Gaussians functions (**Figs. S15-17**), multiplied by the number of spike trimers in each complex and normalized to one. For each spike-antibody, pair the model was globally fit to the three measured concentrations simultaneously. The results of the global fits are shown as red symbols. Background colors corresponds to spike trimer, dimer of trimers and trimer of trimers (orange cyan and pink respectively) and the number of antibodies bound are indicated on the top of each panel. The resultant fitting values for the model parameters are given in **Table S2**.

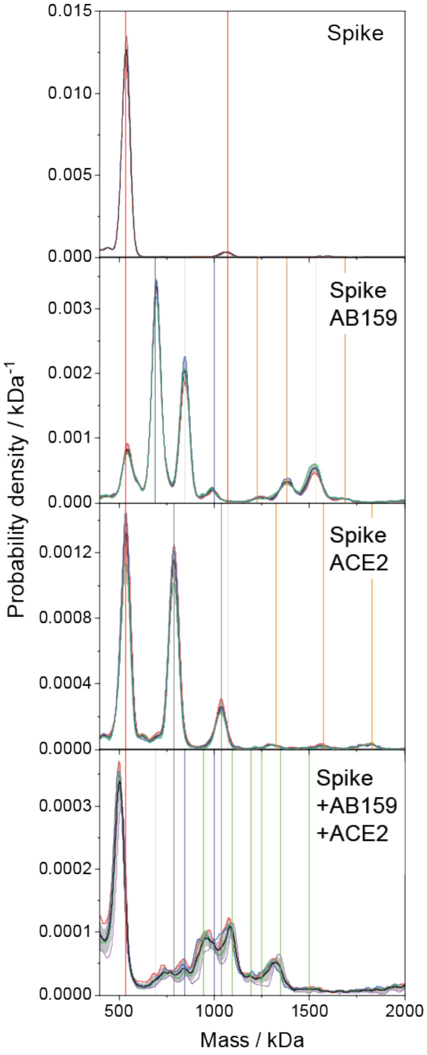


**Fig. S14. Solution MP of ACE2 binding to COVOX159-bound Spike.**

Mass photometry histograms for wtSpike-COVOX159-ACE2 interactions. The top panel shows the spike mass distribution in solution at 17 nM (trimer concentration). The red vertical lines correspond to the spike trimer and dimer of trimers. The second panel shows the histogram of 17 nM spike mixed with 50 nM of COVOX159. The blue vertical lines correspond to the expected spike-antibody masses, and orange lines correspond to spike dimer of trimers bound to 1,2,3 and 4 antibodies. The third panel shows the histogram of 17 nM spike mixed with 100 nM of ACE2, where grey lines correspond to spike trimer bound to 0,1 and 2 ACE2. Orange lines correspond to dimer of trimes bound to 1, 2 and 3 ACE2. The bottom panel shows the mass histogram at a final concentration of 17 nM spike mixed with 50 nM of the COVOX159 antibody and after 10 min of incubation, a final concentration of 70 nM of ACE2 was added to the solution. Four technical repeats were measured following between 5-20 min of additional incubation. Green vertical lines correspond to spike with 1:1, 2:1, 1:2, 3:1, 2:2 and 3:2 159:ACE2 ratios. For all panels, technical repeats are plotted in colour, while black lines and grey error bars represent the average histograms and standard deviations.

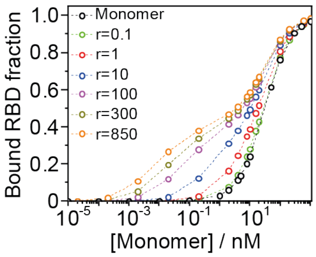


**Fig. S15. Bound RBD fraction as a function of ligand concentration for different surface densities.**

Simulating RBD occupancy as a function of antibody 159 concentrations at different surface densities of spike trimers. Black symbols and dashed line correspond to the monovalent case where the antibody is represented by one Fab domain (no oligomerisation is possible). Colored symbols and lines correspond to the case where binding of a divalent antibody was considered, where oligomerisation is possible. The surface densities of spike in units of particles per μm$^2$ are indicates. Binding curves were calculated by assuming the binding parameters of antibody 159 (**Table S2**).

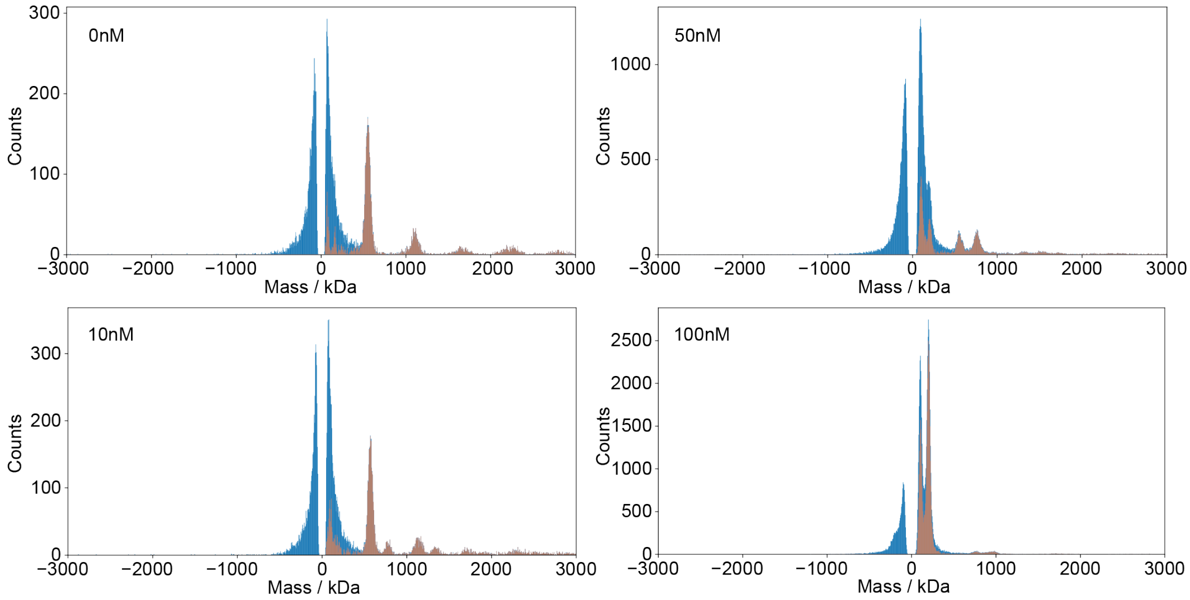


**Fig. S16. Subtraction of low mass impurities (<500 kDa) for rapid dilution experiments.**

Mass histograms when measuring the interaction of spike and ACE2 at µM concentration using the rapid dilution method. Histograms are presented for all mixing concentrations ranging from 0 to 100 nM ACE2 and 50 nM spike. Blue and brown show the raw and subtracted histograms. Subtraction was done by mirroring the unbinding distribution (negative mass) to the positive mass side, and then subtracting the corresponding counts from each bin in the histogram. The decaying distribution is due to small impurities in the spike sumle that increase the noise in the MP movies. Subtracted histograms are also shown in **Fig. 1** and **Fig. S3**. The subtraction does not change the distribution of spike or its interactions with ACE2, but improves the visibility of monomeric and dimeric ACE2 in the low mass range.

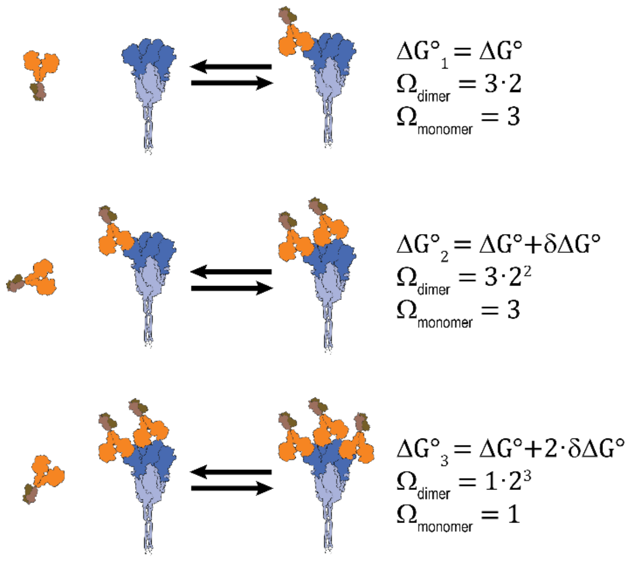


**Fig. S17. Illustration of the interaction model for calculating interaction free energies**

The illustration shows the possible reactions and parameters that were considered for fitting the data presented in Fig. 3d-f. Three possible reactions were considered, for occupying each of the three RBDs on the spike trimer. For omSpike, a maximum of two ACE2 on the same spike were considered since even at high ACE2 concentrations we did not see any significant indication for the existence of three ACE2 dimers bound to a single omSpike. The two fitting parameters for each reaction were the standard free energy change for binding the RBD ($\boldsymbol{\Delta G^{\circ}}$) and a cooperativity free energy factor $\boldsymbol{\delta G^{\circ}}$ that accounts for the possibility that following a previous binding event the subsequent binding is more probable ($\boldsymbol{\delta G^{\circ} < 0}$, positive cooperativity) or less probable ($\boldsymbol{\delta G^{\circ} > 0}$). For each bound state, the degeneracy of the microstates due to the valency of spike and ACE2 was added as $\boldsymbol{\Omega_{monomer/dimer}}$. For the dashed lines in Fig. 3d-f only one parameter was fitted ($\boldsymbol{\Delta G^{\circ}}$) and we set $\boldsymbol{\delta G^{\circ} = 0}$.

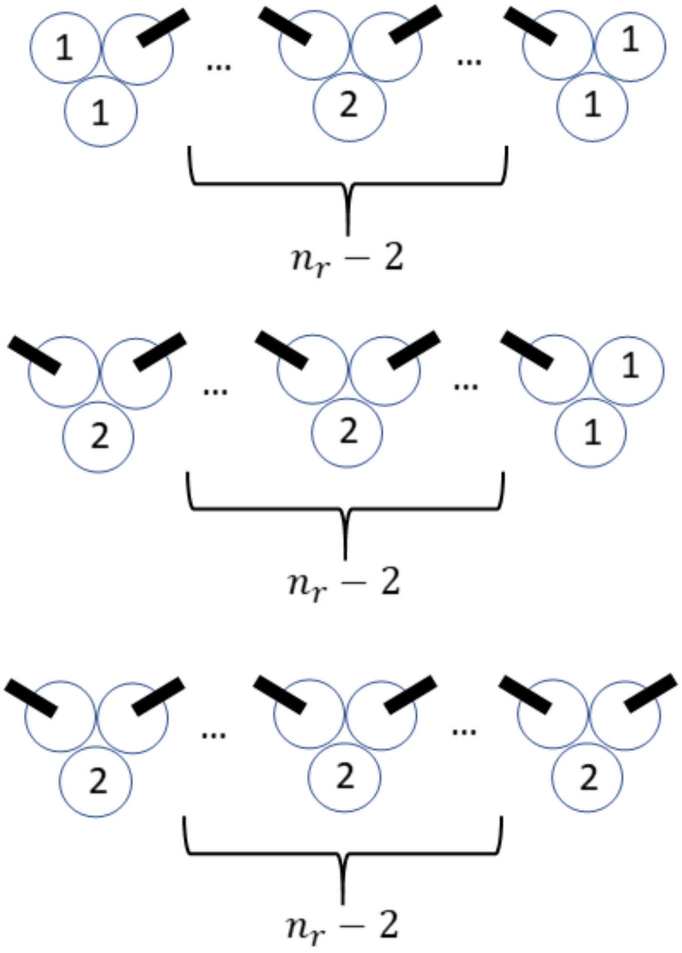


**Fig. S18. Illustration of the distinction between the two spike binding sites used to model the binding of antibody 384 and 150.**

Both antibodies 384 and 150 show negative cooperativity in their binding pattern, therefore, for calculation of the configurational degeneracy we defined different binding states that result in different free energies owing to the binding of different types of sites. Here we show the definition of the two different binding sites (sites 1 and 2). Binding site 1 is defined as a free RBD site that is adjacent to only one prebound RBD, while binding site 2 is a free RBD that is adjacent to two occupied RBD. Owing to the negative cooperativity, the free energy of binding is different for each site type.

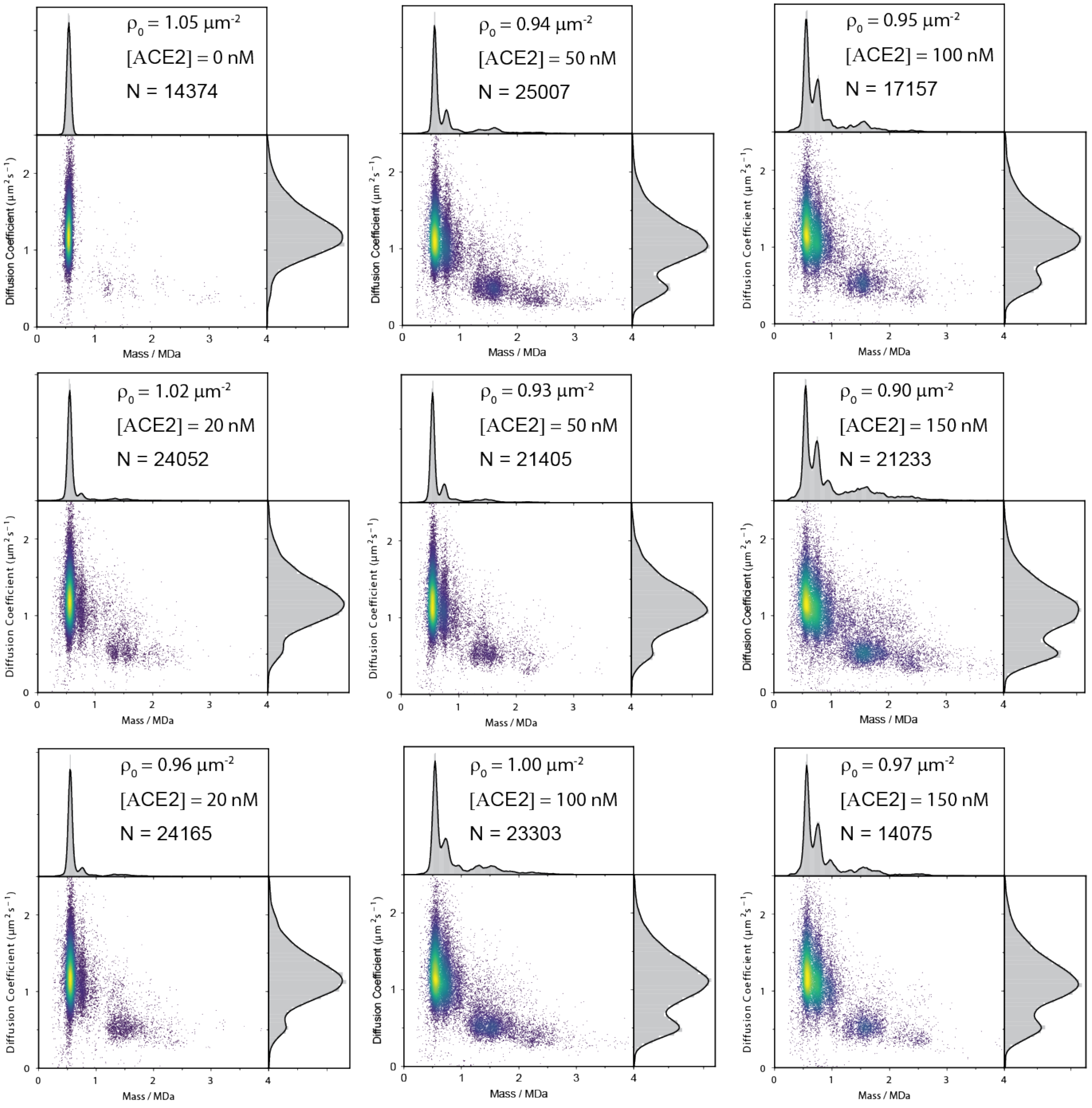


**Fig. S19. 2d plots of omSpike interacting with soluble ACE2**

Two titration data sets of tethered spike at a surface density of ~1 particle/$\mu m^2$ and with increasing concentrations of ACE2.

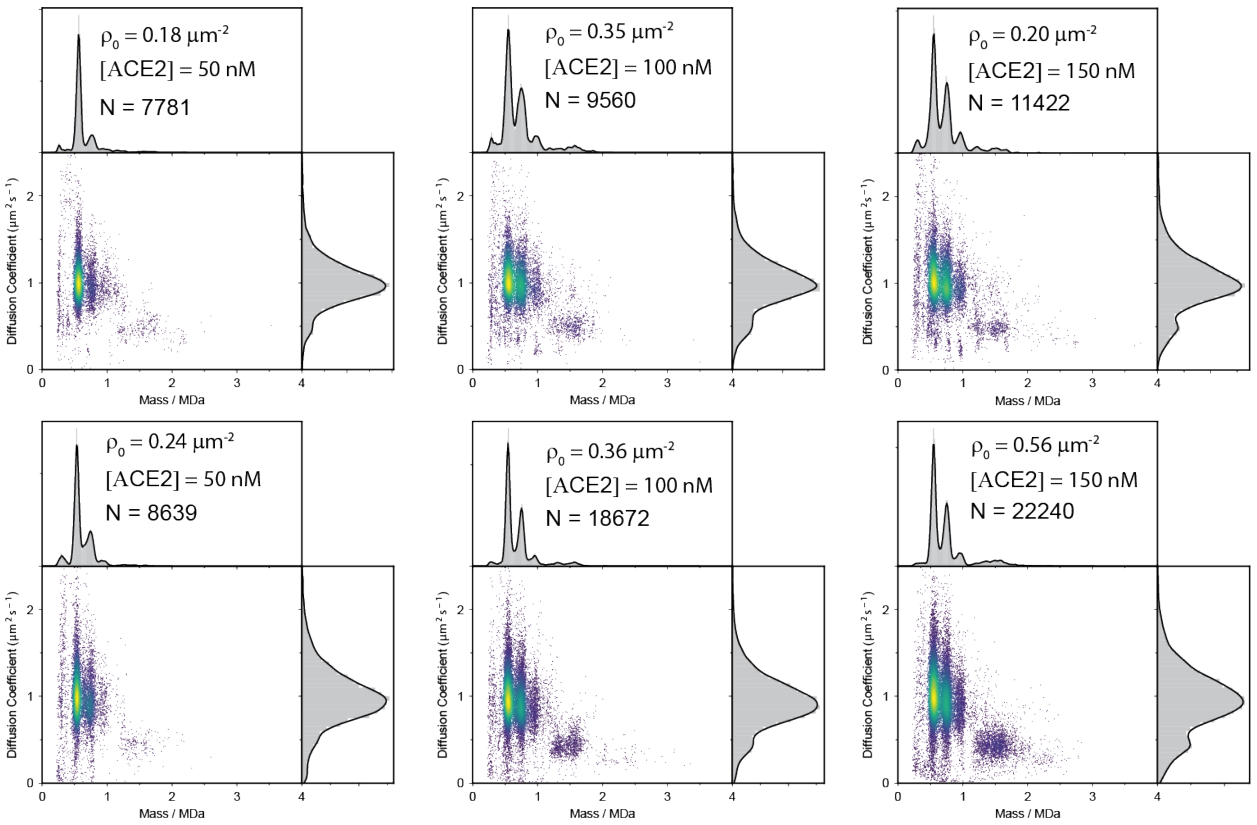


**Fig. S20. omSpike tethered to the supported lipid bilayer interacting with ACE2 as a function of surface density**

Effect of reducing the surface density of spike on ACE2-induced oligomerisation.

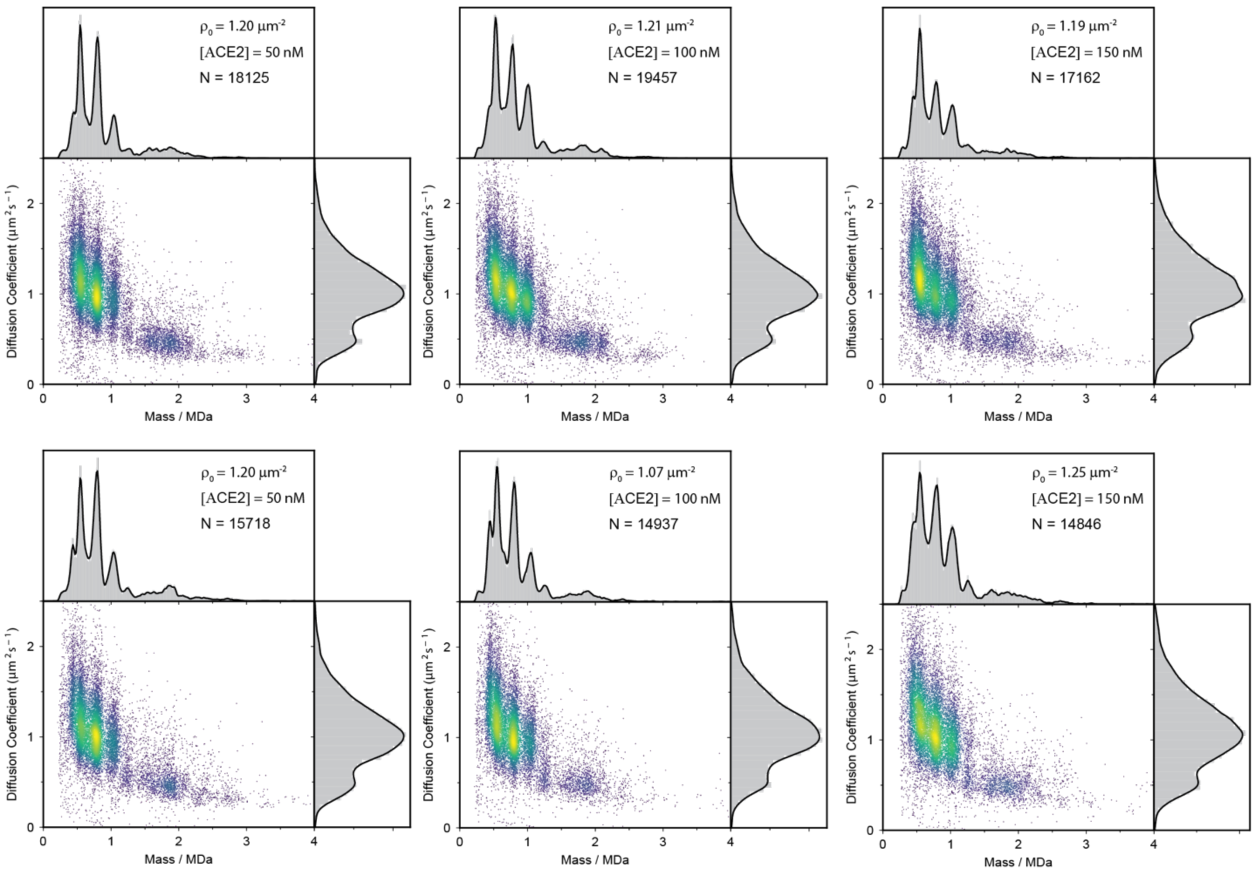


**Fig. S21. 2d plots of wtSpike interacting with soluble ACE2**

Two titration data sets of tethered spike at a surface density of ~1-1.25 particle/μm$^2$ and with increasing concentrations of ACE2. The measured signal at 0 nM of ACE2 iss shown in **Fig. 2**.

| Interaction type | $\Delta G°_{\text{ACE2-ACE2}}$ / $k_B T$ | $\Delta G°_{\text{ACE2-RBD}}$ / $k_B T$ | $K_d^{(1)}$/ nM | $K_d^{(2)}$/ nM | $K_d^{(3)}$/ nM | $K_d^{(4)}$/ nM |
|---|---|---|---|---|---|---|
| ACE2-RBD | -22.3 $\pm$ 0.2 | −21.4 $\pm$ 0.3 | 11.7 $\pm$ 1.6 | 28.2 $\pm$ 9 | 14.1 $\pm$ 2 | 56.4 $\pm$ 18 |
| **Interaction type** | **$\Delta G°_{\text{Spike-ACE2}}$ / $k_B T$** | **$\delta\Delta G°$ / $k_B T$** | **$K_d$ / nM** | | | |
| SpikeWT-mACE2 | -18.51 $\pm$ 0.03 | -0.34 $\pm$ 0.04 | 170 $\pm$ 5 | | | |
| SpikeWT-dACE2 | −19.22 $\pm$ 0.04 | 1.15 $\pm$ 0.06 | 42 $\pm$ 2 | | | |
| SpikeOmi-dACE2 | −18.54 $\pm$ 0.04 | -0.74 $\pm$ 0.07 | 82 $\pm$ 3 | | | |

**Table S1. Summary of the thermodynamic parameters used to model the solution distribution of the interaction between different components of the spike ACE2 interaction.**

| Antibody | $K_d^{(1)}$/ nM | $K_d^{(2)}$/ $\mu m^{-2}$ | $K_d^{(1)}/K_d^{(2)}$/ $nM \cdot \mu m^2$ | $\alpha$ | Δ |
|---|---|---|---|---|---|
| 159 | 5.3 $\pm$ 0.4 | 0.26 $\pm$ 0.04 | 20.55 $\pm$ 3.29 | - | 0.71 $\pm$ 0.02 |
| 150 | 1.3 $\pm$ 0.07 | 2.15 $\pm$ 0.07 | 0.62 $\pm$ 0.05 | -3.01 $\pm$ 0.05 | - |
| 384 | 1.2 $\pm$ 0.04 | 0.11 $\pm$ 0.02 | 11.31 $\pm$ 2.26 | -0.87 $\pm$ 0.06 | 0.67 $\pm$ 0.05 |

**Table S2. Summary of the thermodynamic parameters used to model the distribution of surface complexes between spike and antibodies.**

The parameters correspond to the best fitted model parameters for the 2-dimensional oligomerisation model presented in the Materials and Methods. The statistical errors for the different parameters were calculated by globally fitting different combinations of each data set repeats and concentrations. For example, the data set of antibody 159 contains 3 concentrations and two technical repeats, resulting in 8 different combinations for the global fit. The standard deviation of the fitted parameters from the different combinations was taken as the statistical error.

**Movie S1.**

30 s of a representative ratiometric movie of a solution MP measurement of a mixture of wtSpike at a final concentration of 16.7 nM trimer with an ACE2 final concentration of 100 nM of monomer. The field of view is 3.8×10.7 $\mu m^2$, and the contrast scale is from -0.01 to +0.01. The raw frames were saved at 200 Hz, and the sliding ratiometric processing included a frame averaging of 5 frames. The movie is played back at 30 Hz. The movie was generated using DiscoverMP software v2023R1.2 (Refeyn Ltd., Oxford)

**Movie S2.**

10 s of a representative median subtracted movie of an MP measurement of omSpike tethered to a supported lipid bilayer containing 1% of lipid with a functional NTA(Ni) head group that binds the 6xHis-tag of spike. The field of view is 6.3×9.9 $\mu m^2$, and the contrast scale is from -0.03 to +0.03. The raw frames were saved at 540 Hz with further frame binning of 2 resulting in an effective frame rate of 270 Hz. Median window size was 600 frames. The movie shows 2700 frames between frames 3000 and 5700. The movie is played back at 30 Hz. Scale bar is 1µm.

**Movie S3.**

10 s of a representative median subtracted movie of omSpike, tethered to a supported lipid bilayer containing 1% of lipid with a functional NTA(Ni) head group that binds the 6xHis-tag, following the addition of ACE2 and incubation for 1 h. The field of view is 6.3×9.9 $\mu m^2$, and the contrast scale is from -0.03 to +0.03. The raw frames were saved at 540 Hz with further frame binning of 2 resulting in an effective frame rate of 270 Hz. Median window size was 600 frames. The movie shows 2700 frames between frames 6000 and 8700. The movie is played back at 30 Hz. Scale bar is 1µm.

**Movie S4.**

10 s of a representative median subtracted movie of HexaPro stabilised wtSpike, tethered to a supported lipid bilayer containing 1% of lipid with a functional NTA(Ni) head group that binds the 6xHis-tag. The field of view is 6.3×9.9 $\mu m^2$, and the contrast scale is from -0.03 to +0.03. The raw frames were saved at 540 Hz with further frame binning of 2 resulting in an effective frame rate of 270 Hz. Median window size was 600 frames. The movie shows 2700 frames between frames 301 and 3000. The movie is played back at 30 Hz. Scale bar is 1µm.

**Movie S5.**

10 s of a representative median subtracted movie of HexaPro stabilised wtSpike, tethered to a supported lipid bilayer containing 1% of lipid with a functional NTA(Ni) head group that binds the 6xHis-tag, following the addition of 5 nM of COVOX384 and incubation for about 1 h. The field of view is 6.3×9.9 $\mu m^2$, and the contrast scale is from -0.03 to +0.03. The raw frames were saved at 540 Hz with further frame binning of 2 resulting in an effective frame rate of 270 Hz. Median window size was 600 frames. The movie shows 2700 frames between frames 301 and 3000. The movie is played back at 30 Hz. Scale bar is 1µm.
